# Historical adaptation to transposable element invasions in *Drosophila* is primarily due to insertions in existing piRNA clusters

**DOI:** 10.64898/2026.11.09.764865

**Authors:** Adriana Ludwig, Joanne S. Griffin, Andrea J. Betancourt

## Abstract

Transposable elements (TEs) are genomic parasites that shift between host species, which may allow them to escape sequence-specific host silencing mechanisms. One such silencing mechanism, the piRNA pathway, is the main line of defense in the germline of animals, and acts through small RNAs targeting specific TE sequences.

Though many mechanistic details of this pathway are known, how it evolves to respond to a new transposable element invasion is not. Here, we examine the piRNA-based response to three recently invading TEs in *Drosophila simulans,* the *P-element, Shellder and Spoink*. To this end, we assembled genomes of 22 strains, annotated piRNA-producing loci (piRNA clusters) in each strain, and located insertions of the new TEs within these loci. We find that piRNA clusters are highly polymorphic among strains and piRNAs targeting the new TEs come mostly from clusters that likely pre-date the invasions, though there is also some evidence of piRNAs generated *de novo*, particularly for the long-terminal repeat (LTR) elements. Moreover, resistance to each TE appears to have evolved mostly independently in the different strains through distinct insertions, often in different clusters, although some loci, such as *flamenco* for *Shellder,* are recurrent targets.

## Introduction

Transposable elements (TEs) are the canonical selfish genes; they replicate by increasing their numbers in genomes, regardless of harm done to their hosts [1–5]. This evolutionary strategy is undeniably successful— TEs are ancient, taxonomically widespread, and they make up substantial proportions of most eukaryotic genomes [6, 7].

As a consequence, TEs have a profound impact on host genome evolution and fitness [8–10]. TE sequences, proteins, and replication machinery have each been co-opted numerous times by their hosts, contributing to evolutionary novelty [11–13]. Conversely, TE replication can be harmful to the host [reviewed in [14, 15]]. These harmful effects can include insertion mutations [16, 17], genome rearrangements [18, 19], epimutations [20, 21] and sterility [22, 23]. Newly transferred, unsuppressed TEs with high replication rates are likely to be especially harmful, and impose strong selection on their hosts to evolve suppression rapidly [17], particularly when they cause lethality or sterility in offspring [1]. In fact, newly invading transposable elements can rapidly become inactive or suppressed, as has been repeatedly shown in the laboratory [24–27].

Once a TE family is suppressed to the extent that it no longer replicates within genomes, it is effectively evolutionarily dead [28, 29]. Genomes are littered with these inactive TE sequences, which are typically incomplete or otherwise degraded [30, 31]. Ultimately, shifting among susceptible host species may be key to the long-term evolutionary persistence of TEs [32, 33]. This is because newly transferred TEs likely enjoy a brief period where they escape host silencing and can transpose relatively freely. In fact, TEs are well-known to transgress species boundaries in eukaryotes as well as prokaryotes, with bacteria, large viruses and large TEs implicated as potential means of transfer, possibly with the assistance of eukaryotic vectors [34–38].

Horizontal transfer of TEs between eukaryotic species was originally discovered in *Drosophila melanogaster*: In that species, a DNA transposon, the P-element, is present in strains collected after about 1970, but absent from strains collected prior to that time, and from its close relatives [39, 40]. A very similar P-element was found in a distantly related species, implying horizontal transfer without hybridisation [41]. The lack of suppression in naïve flies likely enabled its rapid spread in both *D. melanogaster* and later, in the 21^st^ century, *D. simulans* [23, 42, 43].

In general, new TEs may escape suppression because many, though not all, host silencing mechanisms recognise TEs by their sequences [44]—*e.g.*, Krab-associated zinc-finger proteins (KZFPs) which recognise specific sequence motifs [45, 46], or small RNAs cognate to particular TE sequences [47]; new TE families may have sequences that are unrecognised by these mechanisms. Other silencing mechanisms, though not sequence-specific, would be equally ineffective against very new TE families with few copies in the genome. Repeat-induced point mutation in fungi, for example, requires multiple copies of a TE family to be present [48–50].

Eventually, however, hosts do evolve suppression. In most animals, suppression of transposition in the germline is mediated partly through maternally expressed piRNAs, small RNAs associated with Piwi-clade Argonaute proteins [51, 52]. Host piRNAs are produced from genomic source loci known as piRNA clusters. These can be either large dedicated loci containing many TE sequences or individual TE insertions, and piRNA production can also be stimulated by nearby piRNA-producing sequences [53–57]. These clusters can evolve by acquiring new TE sequences, and new piRNA clusters can arise in fly genomes, possibly spreading from foci centred on individual TE copies [58, 59].

Suppression of a new TE can therefore potentially occur through multiple routes: established piRNA clusters may serve as traps, ensnaring TEs that transpose into them [60–62], or, *de novo* piRNA clusters can form, through mechanisms that are not entirely clear, but which may involve antisense TE transcripts [54, 56, 57]. Laboratory experiments showing suppression of the P-element evolving in the laboratory suggest that production of P-element cognate piRNAs is initiated on a rapid time-scale, during course of the experiment [27, 63]. In one case, these appear to originate from a *de novo* piRNA cluster [63]. However, there are few empirical tests showing how TE silencing evolves in natural populations.

Here, we investigate how silencing of newly invading TEs is established, using *D. simulans* to provide a long overdue empirical example of the evolution of TE silencing in natural populations. *D. simulans,* the sister species of *D. melanogaster,* was invaded by P-element only recently, such that many strains collected prior to the invasion are still in culture [23, 64]. About a decade earlier, *D. simulans* was also invaded by two LTR-retrotransposons, *Shellder* and *Spoink* [65, 66], providing two additional cases of recent invasion. We leverage long-read and small RNA sequencing to annotate piRNA source loci in uninvaded and invaded *D. simulans* strains and to understand how piRNA silencing evolved in this species. As found previously [58, 59], the piRNA source loci in our strains range from regions as small as a single copy of a TE to large genomic regions. Some source loci are fixed (i.e., present in all strains), while others show presence/absence polymorphism. We show that, our *D. simulans* strains are protected from the newly invading TEs due to piRNAs produced at long-established piRNA clusters, largely, though not exclusively, supporting the trap model of adaptation to TE invasions [61, 67].

## Materials and Methods

### Fly strains, DNA extractions and Nanopore sequencing

We obtained 22 *D. simulans* strains from both immediately before (12 strains) and after (10 strains) the invasion of this species by the P-element, isolated HMW DNA and sequenced each line using Oxford Nanopore ligation sequencing kits run on a GridION (Centre of Genomic Research, Liverpool) for 72 hours.

### Small RNA extraction and sequencing

Small RNA sequencing libraries were prepared from RNA extracted from dissected ovaries of 3-5 day-old females using the NEBNext small RNA library prep kit, including an additional 2S rRNA blocking step [68], and sequenced on one lane of the Illumina NovaSeq X Plus using 10B chemistry at the Centre for Genomic Research in Liverpool.

### Genome assembly and scaffolding

After base-calling, trimming and filtering, reads were *de novo* assembled using Flye v2.9.2 [69], and assemblies polished with Flye, Racon v1.5.0 [70] and Medaka v1.11.3 (https://github.com/nanoporetech/medaka). For four strains, Illumina reads were available, and one round of polishing was done with Pilon v1.24 [71]. Craq v1.0.9-0 [72] was used to detect structural errors and break chimeric contigs, and Kraken2 v2.1.3 [73] was then used to identify and eliminate contigs from potential contaminant sources. We used RagTag v2.1.0 [74] to scaffold the resulting filtered contigs on the Prin_Dsim_3.1 (GCF_016746395.2) reference genome, and assessed the quality of the final assembly with BUSCO v5.7.1 [75], Quast v5.2.0 [76] and Craq v1.0.9-0 [72].

### Pangenome alignment

We generated a pangenome alignment using Cactus v2.9.7 [77], implemented via Singularity, with Prin_Dsim_3.1 as the reference genome. Before alignment, each genome was softmasked using maskfasta from bedtools v2.30.0 [78] and its respective TE annotation generated with a preliminary TE library.

### TE library curation

We generated two TE libraries, a fully manually curated *D. simulans*-specific library and a *Drosophila* library. The *D. simulans* library was obtained using a workflow combining RepeatModeler v2.0.3 [79], MCHelper v1.7.0 [80] and TE-trimmer v1.4.0 [81] applied successively to the *D. simulans* reference genome (Prin_Dsim_3.1) and then to each of 22 genomes assembled here. Additional sequences were added using Pantera after pangenome alignment [82]. Redundancy was eliminated by clustering potential TEs using the 80/80 rule [83]. Consensus sequences were inspected using TE-aid (https://github.com/clemgoub/TE-Aid) and manually curated following [84]. The final *D. simulans* library consists of 224 fully manually curated consensus sequences.

### A combined Drosophila

TE library was obtained by adding to our *D. simulans* library 157 curated TE sequences from *D. melanogaster* [85], supplemented with 5 additional sequences from Repbase23.11 [86] and the BDGP TE library [30], and 2,263 from diverse *Drosophila* species [87], inspected to eliminate satellites, rDNA, low-complexity sequences, and clear chimaeras or non-TEs. The final combined TE library comprises 2,649 sequences with some redundancy.

### TE annotation of genome assemblies

We identified the locations of TE sequences in our genome assemblies with a pipeline that combines RepeatMasker v 4.1.1 [88] and blast [89]. The curated *D. simulans* TE library was used in the first module to annotate TE copies with high within-family sequence similarity, and the combined *Drosophila* library was used later to annotate divergent TE fragments. We estimated the number of TE copies using “One code to find them all” [90], a set of scripts to parse RepeatMasker results obtained with the curated *D. simulans* TE library.

### piRNA cluster annotation of genome assemblies

We used evidence from RNAseq of small RNAs to annotate piRNA clusters for each strain. Post-trimming with Cutadapt v1.2.1 [91], we filtered for lengths of 23-29 nucleotides, eliminated reads mapping to annotated rRNA, tRNAs, miRNA and miscRNAs [92], and to the sense strand of genes (except for those reads matching TE sequence). TBr2_collapse and TBr2_duster from the NGS toolbox [93] were used to collapse the remaining reads, retaining only unique piRNAs and removing low-complexity reads. We used the filtered reads to perform two complementary sets of annotations on each of our 22 assemblies broadly following [59], using Protrac [93], and a coverage-based approach. To perform the coverage-based annotation, we used unique mapping and a maximum of 1 mismatch, retaining regions that met the criteria of sufficient coverage (≥25 RPKM), 1U bias >60%, overlapping regions containing TEs, and not overlapping gene sequences. Predicted piRNA clusters were first merged within each strain across replicates using a 10-kb window. Clusters were filtered to remove gene-overlapping regions following [59] and those lacking TE content or shorter than 2 kb. The coordinates of the predicted clusters were transferred to the reference genome using halLiftover [94], and shared clusters across strains were identified by merging lifted coordinates with bedtools.

### Ping-pong signal and sense/antisense piRNA expression

Filtered reads were mapped to the curated *D. simulans* TE library using Bowtie v1.3.1 [92], allowing at most 2 mismatches. To quantify the ping-pong signature, we used the Python script signature.py (https://github.com/ARTbio/tools-artbio/blob/main/tools/small_rna_signatures/signature.py), using the method originally described by [95]. A ping-pong signal was considered significant when the 10-nucleotide overlap had at least 10 supporting read pairs and a Z-score ≥ 2. Sense and antisense TE counts were normalized by TE length and by the total number of mapped miRNA, producing miRNA-normalized RPKM values for each TE. *D. simulans* miRNA were obtained from miRbase [96].

### Detection of unassembled TE copies from long reads

Filtered long reads from the 22 strains were mapped to the Prin_Dsim_3.1 reference genome (GCF_016746395.2) using [97] and Sniffles2 [98] was used to call structural variants from the alignments. Custom Python and Bash scripts were used to extract insertion sequences, identify P-element, *Shellder*, and *Spoink* insertions via blastn, and extract the TSD sequences associated with each insertion. These insertions were intersected with the piRNA cluster annotations. Insertions were assigned to a piRNA cluster if they fell within a region annotated as piRNA cluster in any strain.

A detailed description of Materials and Methods is available as Supplementary Text and as a GitHub repository (https://github.com/adriludwig/Drosophila_simulans_piRNA_clusters).

## Results

### Genome assembly, TE curation and annotation

We sequenced and assembled genomes for 22 *D. simulans* lines collected in the years between 2002 and 2024 from various locations worldwide (Table S1), using long reads with an average coverage of 109-fold (range 25-201). Initial assemblies exhibited moderate to high contiguity (median N50 = 14.4 Mb), which was further improved by scaffolding the contigs on a reference genome (Fig. 1A). N50 values exceeded 20 Mb across all samples, close to the length of chromosome arms for this species, 21/22 genomes had S-AQI scores >90% (Table S2), and the assemblies capture most of the protein-coding gene content as measured by BUSCO scores (Fig. S1). Potential misassembles are given in Table S3.

**Figure 1.**
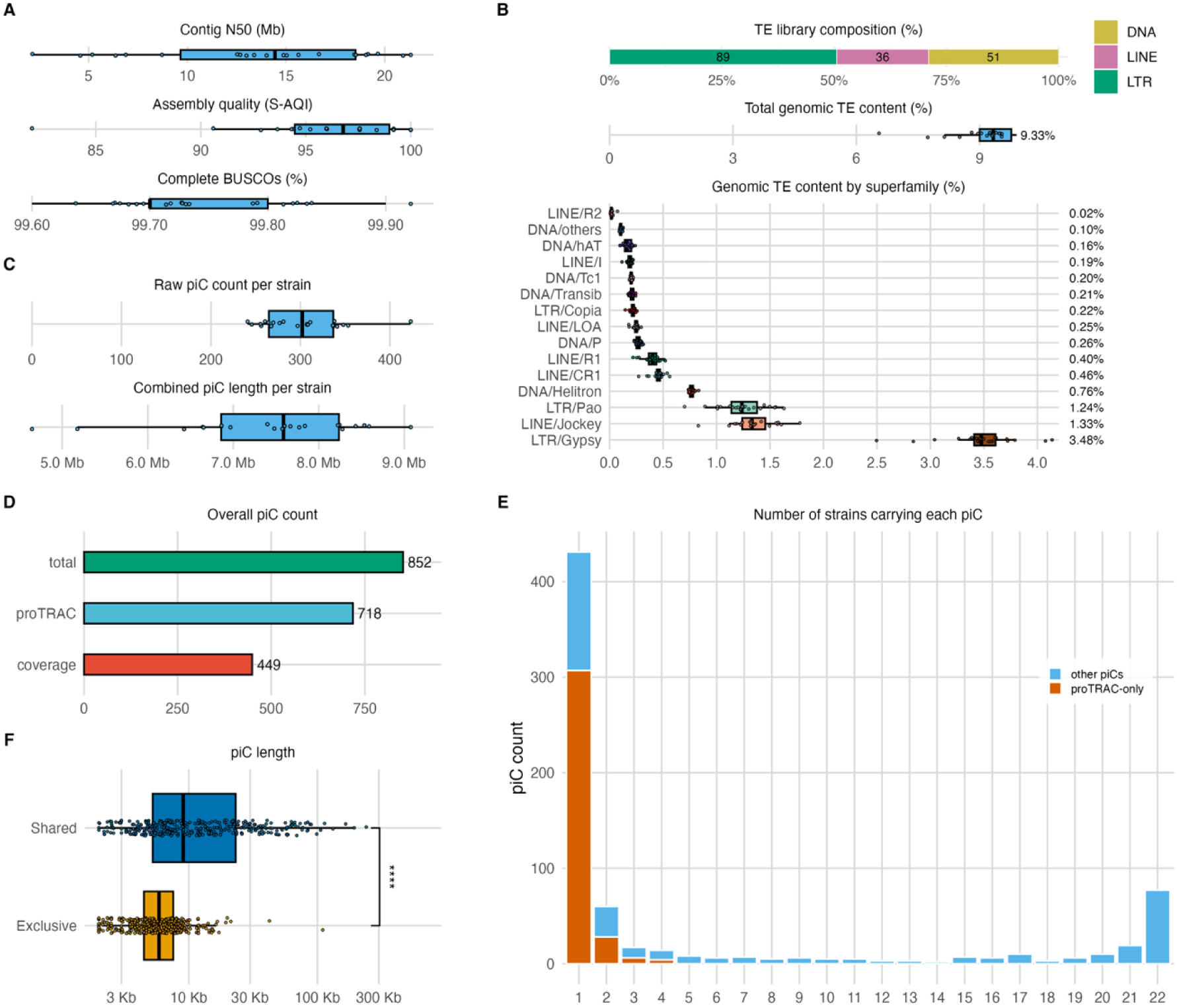
Assembly quality, TE composition, and piRNA clusters (piCs). (**A**) Metrics for genome assembly quality across various strains. Boxplots illustrate contig N50 (Mb), S-AQI (a structural assembly quality index derived from CRAQ based on normalised counts of clip-based structural errors per Mb), and the percentage of complete BUSCOs. (**B**) Top panel: The contribution of major TE groups (DNA, LINE, LTR) within the curated TE library is shown as percentages, with the number of consensus sequences indicated on the stacked bars. Only complete or nearly complete TE consensus sequences were counted (176 sequences). Mid panel: Total TE genomic contribution across strains, with median percentage displayed on the right. Bottom panel: Genomic contribution of TE superfamilies across strains, with median percentages displayed on the right. TE content was calculated per strain as the percentage of the assembled genome length occupied by TE annotations. (**C**) Number of piRNA clusters detected per strain and combined piRNA cluster length in Mb. **(D)** Overall number of non-redundant piRNA clusters identified across the 22 strains, together with the number detected by proTRAC or the coverage-based approach in at least one strain. **(E)** Distribution of piRNA clusters according to the number of strains in which they are detected. Only clusters in which the syntenic regions were represented in all genome assemblies were considered (710 of 852 clusters). Clusters predicted by proTRAC-only are shown in a different colour from the others (predicted by coverage-only or both). (**F**) Comparison of piC length between shared (at least 2 strains) and exclusive (only one strain) clusters annotated with proTRAC and/or coverage-based methods, including all 852 clusters (****Wilcoxon rank-sum test, p = 6.2 x 10^-30^).

To facilitate annotation of the TE content in our assemblies, we first obtained a comprehensive, manually curated library of TEs for *D. simulans*. The final non-redundant library of consensus sequences consists of 224 sequences, with 176 corresponding to complete or nearly complete TEs, roughly half of which are LTR elements (Fig. 1B). Several of these families show high sequence similarity to *D. melanogaster* TE families (Table S4), reflecting shared ancestry and/or ongoing horizontal transfer [99, 100]. The mean TE content across strains in the genome assemblies was 9.3% (range 6.6% - 10.9%; Fig.1*B*)—almost certainly an underestimate, as the unassembled regions are expected to be enriched for repetitive elements. Accordingly, there is a correlation between genome assembly quality and TE content (Fig. S2), consistent with better assemblies capturing more repetitive DNA. The TE content in the assemblies varies by superfamilies, with the highest abundance (as measured by bp coverage) being LTR/Gypsy, LINE/Jockey, LTR/Pao and DNA/Helitron. These four superfamilies are also the most represented in terms of copy number (Fig. S3), roughly consistent with previous estimates of TE content in *D. simulans* [101, 102].

We also see signatures of changing TE content through time, as expected. The P-element was absent from 13 strains (Fig. 2; Table S1), which were collected before the P-element became common in 2014, including the reference genome [23, 103]. Two other TEs, *Spoink* and *Shellder*, are absent in the reference genome but present in all other strains; these TEs invaded *D. simulans* in the 1990s [65, 66]. In addition, one element (Mariner3_Ds/Mos-1) is at low copy number in most strains (mean copy number 1.63), and absent from two (Fig. S4), consistent with presence/absence polymorphism previously reported for this family in natural populations [104].

**Figure 2.**
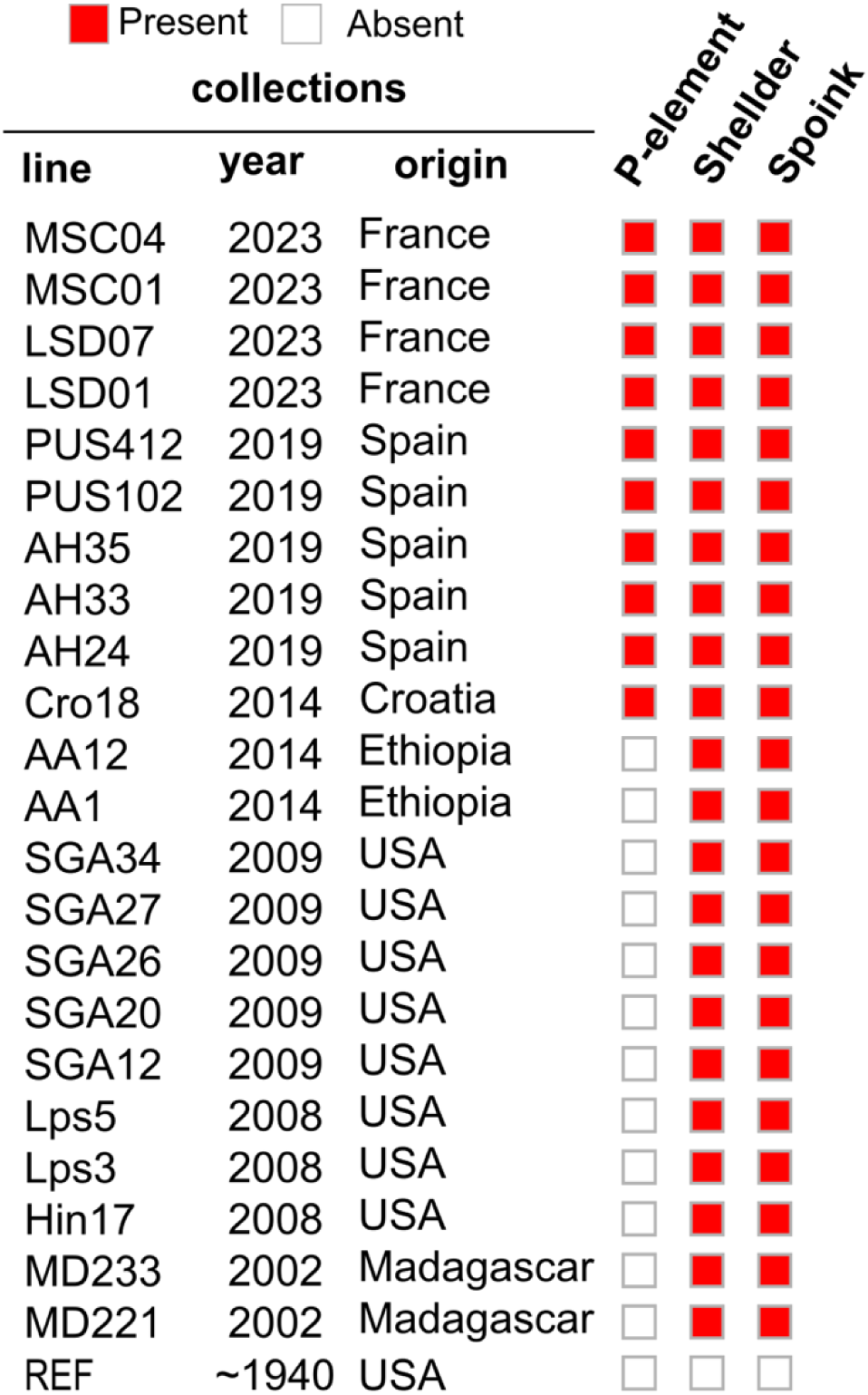
Distribution of P-element, *Shellder*, and *Spoink* across lines. Presence (red) and absence (empty) of each element are shown for strains grouped by collection year and origin. REF strain corresponds to Prin_Dsim_3.1 (GCA_016746395.2).

### piRNA cluster annotation for D. simulans

We annotated source loci for piRNAs, or piRNA clusters [60], for each strain separately (2-6 small RNA library replicates, Table S1), using two complementary methods: proTRAC [93], which employs a probabilistic algorithm to identify these clusters through multimapping, and a coverage-based approach, which identifies clusters based on the density of uniquely mapped reads. Our annotated piRNA clusters are concentrated in pericentromeric regions, mainly those of chromosomes 2 and X, which have higher density and larger clusters (>10kb), qualitatively similar to patterns seen in *D. melanogaster* [59, 105]; though the third chromosome also has a high density of clusters in its pericentromeric region, which are fragmented between 3L and an unplaced contig (Fig. S5).

Each strain had hundreds of predicted clusters (mean = 301; range 241 – 423; Fig. 1C), though the exact numbers are influenced by the decision to split clusters at genes (following [59]. Our cluster annotations are reproducible; on average, 88.6% of all clusters identified in a strain are shared across replicates (Fig. S6).

The annotated clusters comprised large fractions of the assembled genomes, ∼7.4 Mb, or about 5.8% of our assemblies (range 4.6 - 9Mb; 3.8%-6.9%; Fig. 1C; Fig. S7). This overall substantial piRNA cluster content is consistent with previous annotations of piRNA clusters in *D. melanogaster* [59] and somewhat lower than that recently reported for *D. simulans* (13.75-16.35 Mb) [58], likely reflecting differences in annotation criteria.

### piRNA cluster polymorphism in D. simulans

We then compared piRNA cluster presence/absence polymorphism across strains. As piRNA clusters show high within-cluster sequence turnover [58, 59, 67, 106], we identified common clusters by synteny rather than within-cluster sequence similarity. To this end, we lifted the coordinates of the clusters from each strain to the reference genome and intersected them to find piRNA clusters with overlapping reference locations. We were able to identify reference genome locations for most of the annotated clusters in each strain (mean = 232; range 186-301). An average of 42 clusters per strain could not be transferred, comparable to the 30 reported in a similar analysis in *D. melanogaster*, which has a more contiguous reference genome [59]; accordingly, non-lifted clusters were disproportionately found on raw, unscaffolded contigs (52.8%). This cross-strain intersection revealed a total of 852 unique regions, representing the non-redundant repertoire of predicted piRNA clusters across the 22 strains (Fig. 1D; Table S5). Of these, 718 (84.3%) were supported by proTRAC and 449 (52.7%) by the coverage-based method in at least one strain (Fig. 1D). These clusters are largely composed of TE sequences, as expected, with a mean TE sequence coverage of 77.8% across all strains (SD = 21.8%).

We next examined the cluster occupancy frequency, i.e. the number of strains in which each cluster is present. Because the absence of a cluster prediction in a strain may reflect a missing genomic region rather than a true absence, we restricted this analysis to clusters located in regions present in all 22 strains (*n* = 710 clusters). The resulting frequency distribution is asymmetrically U-shaped (Fig. 1E; Fig. S8), with most clusters being strain-specific (431, or 60.7%) or shared across all strains (77, or 10.8%). This skewed distribution is reminiscent of a site-frequency spectrum for alleles with low mutation rates and a degree of purifying selection [107]. However, around 71% of the exclusive clusters (307/431) were predicted only by proTRAC. Because very recent TE insertions have not yet accumulated sufficient sequence divergence, piRNAs derived from them are unlikely to map uniquely, making the multi-mapping approaches used by proTRAC particularly important for detecting *de novo*, low-frequency piRNA clusters. Notably, approximately 80% of exclusive clusters consist of a single TE insertion from active LTR or LINE elements (*e.g. Roo*, *Doc*, *412*, *I*, *Shellder*, *Doc6*, and *Max*), supporting the interpretation that at least a subset might correspond to *bona fide* piRNA source loci recently established by TE insertions. Nevertheless, because multi-mapping reads can be misassigned, some of these low frequency clusters may also be false positives.

Annotated shared clusters are significantly longer than exclusive ones (mean shared 19.7 kb vs. exclusive 6.5kb; Wilcoxon rank-sum test, *p* = 6.2e-30 Fig. 1F) and mean cluster size was positively correlated with occupancy (Pearson’s *r* = 0.59, *p* = 4.8e-80⁻^80^; Fig. S9). This size-occupancy relationship aligns with previous observations in *D. melanogaster* [59] and *D. simulans* [58], consistent with a dynamic ’birth-and-death’ model of piRNA cluster evolution, in which most new, small clusters are rapidly lost, while a select subset persists, increases in population frequency, and expands through trapping new TE insertions.

### Most P-strains have P-element insertions in established clusters whose existence pre-dates invasion

To investigate piRNA-based silencing of the P-element, we examined both P-element piRNA production and the genomic context of predicted piRNA clusters harboring P-element copies.

We find that all ten strains containing P-elements produce both sense and antisense piRNA cognate to the P-element sequence (Fig. 3A,B; Fig. S10). These piRNAs exhibit the 10-nucleotide overlap enrichment (minimum pair counts = 10; Z-score >=2; Fig. 3A, Fig. S11) characteristic of ping-pong amplification as expected for a germline-expressed TE [108]. In contrast, strains lacking a P-element show no detectable P-element-derived piRNAs (Fig. 3B, Fig. S10).

**Figure 3.**
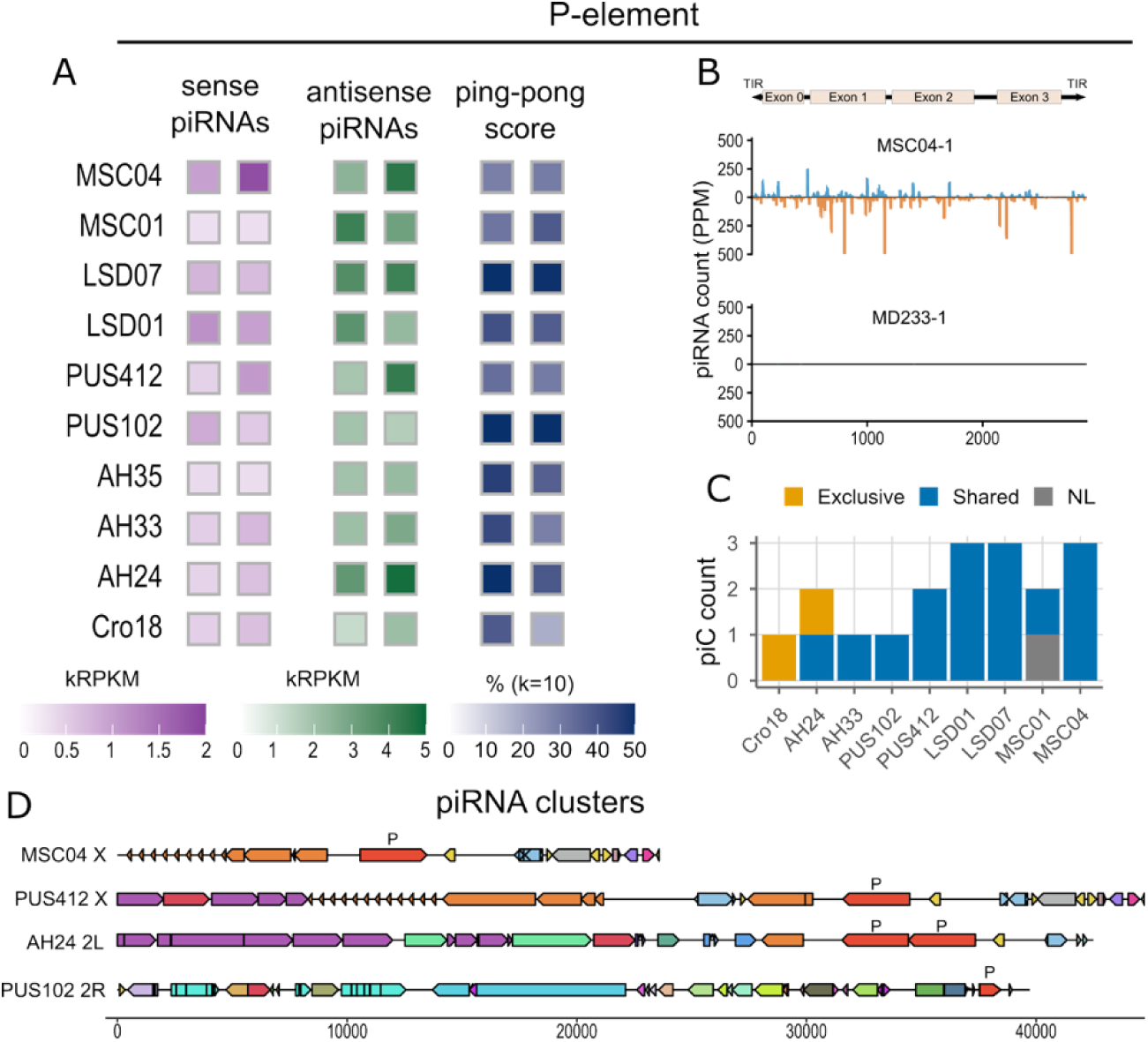
P-element defence. **(A)** Heatmaps of sense and antisense piRNA levels mapping to the P-element are shown in kRPKM (thousand reads per kb per million mapped miRNA) and the ping-pong scores (fraction of 5′-to-5′ sense/antisense piRNA pairs that overlap by exactly 10 nucleotides out of all overlaps) for strains containing the P-element. Two biological replicates were plotted per strain, with random sampling when more than two libraries were available. (**B**) Representative coverage profiles for two strains: MSC04 contains the P-element and shows sense and antisense piRNAs (in PPM) mapping along the P-element sequence, whereas MD233 lacks the P-element and shows no mapping. (**C**) Number of piRNA clusters harbouring the P-element per strain, colored by status (Exclusive, Shared, NL – non-lifted); note that several clusters harbour P-elements in multiple strains, and are represented multiple times (*n*=14 independent clusters). (**D**) Illustration of piRNA clusters having the P-element, with colored segments indicating distinct TEs and orientation (see Fig.S13 for full legend).

We next asked which model of piRNA production for a newly acquired TE is more consistent with our data, the trap model [61, 67] or the *de* novo model [56, 57]. The trap model predicts that most P-element source loci are due to P-element insertions in piRNA clusters that were already present before the P-element invasion. The *de novo* model predicts that P-element source loci are primarily located outside of these established piRNA clusters, perhaps from isolated TE sequences. To distinguish between these, we used a strategy that does not rely on assigning piRNAs to specific P-element copies, for two reasons. For one, P-element copies are highly similar, and uniquely assigning small RNA reads to a copy is often not possible. For another, piRNAs generated from secondary biogenesis may originate from transcripts of any P-element copy [60]. Instead, we assumed that clusters that occur in most or all the uninvaded strains were already established prior to the P-element invasion; we refer to these as ‘established’ clusters. Our test is necessarily one sided—we can confidently identify strains where invading TEs are inserted into established clusters. For strains where no such insertions are identified, our test is inconclusive, as piRNAs may be produced by established clusters not included in our assemblies, and/or from isolated TE insertions that are potential *de novo* clusters.

Of the ten P-element-carrying strains, eight have P-element insertions in at least one established cluster (all but Cro18 and AH35), with insertions in 11 distinct clusters (Fig. 3C, Table S6). Except for one insertion shared between two strains, these insertions are independent, suggesting that resistance to the P-element evolved multiple times through repeated insertions into regions that were already established as piRNA clusters prior to the invasion. For the two exceptional strains, there do not appear to be P-element insertions in pre-existing clusters. Instead, P-element cognate piRNA production seems to originate from source loci elsewhere throughout the genome. In AH35, no P-element containing cluster was predicted despite P-element cognate piRNA production; we cannot exclude that such a cluster exists in an unassembled region of the genome. In Cro18, the primary source of P-element piRNAs appears to be an exclusive, likely a *de novo* cluster (supported by reads mapping to unique flanking sequence) in which a P-element is inserted antisense to a protein-coding gene (LOC6724712) and sense to an overlapping uncharacterized lncRNA (LOC120285219; Fig. S12).

One particular region, a ∼41 kb region near the telomere of the X chromosome, appears to be a fixed piRNA cluster (present in 21/21 strains with sequence in the region), and has an unusual density of P-element insertions: four insertions in three strains. Three of these (in MSC04, LSD07, and PUS412) lie less than 150 bp apart. Despite their proximity, these insertions are distinct transposition events, as evidenced by their different orientations and distinct target site duplications (TSDs). MSC04 contains an additional insertion ∼1.5 kb away. A similar 87.5 kb telomeric cluster on chromosome 2L contains two nested P-element insertions shared between two strains (AH24 and AH33) and an independent insertion in MSC04, ∼ 170 bp apart from this site. In fact, there is a high similarity between the telomeric clusters on these two chromosomes, making it difficult to identify the exact location of these insertions, as the P-element is embedded within the same TE fragments in both regions (Fig. 3D, Fig. S13).

### piRNA production for other recent invaders of D. simulans, Shellder and Spoink

We extended our analysis to include *Shellder* and *Spoink*, two other invaders that recently spread in *D. simulans* [65]. Both elements showed abundant cognate piRNAs in all of our strains (Fig. 4A; Fig. S14; Fig. S15). We asked whether these piRNAs correspond to both strands of the element, as this may shed light on the expression niches of these TEs and their cognate piRNA sources. In fact, these might be expected to differ: while these elements share a similar overall structure, with *gag* and *pol* genes typical of LTR retrotransposons, *Shellder* additionally encodes an envelope (*env*) gene. This distinction is relevant because *env*-containing LTR elements can exhibit retrovirus-like properties and, rather than being directly expressed in germline cells, these elements can be expressed in somatic follicle cells, from which they can enter germ cells [109–113]. Consistent with these biological differences, the two types of LTR elements are controlled by separate piRNA pathways. Retrovirus-like elements are silenced in somatic cells by piRNAs generated from unistrand clusters, which primarily produce piRNAs antisense to the TE transcripts and lack ping-pong amplification [108, 114, 115]. In contrast, germline elements, including LTRs lacking an *env* gene, are usually silenced by dual-strand clusters expressed in the germline, where their piRNAs are amplified through ping-pong amplification [60].

**Figure 4.**
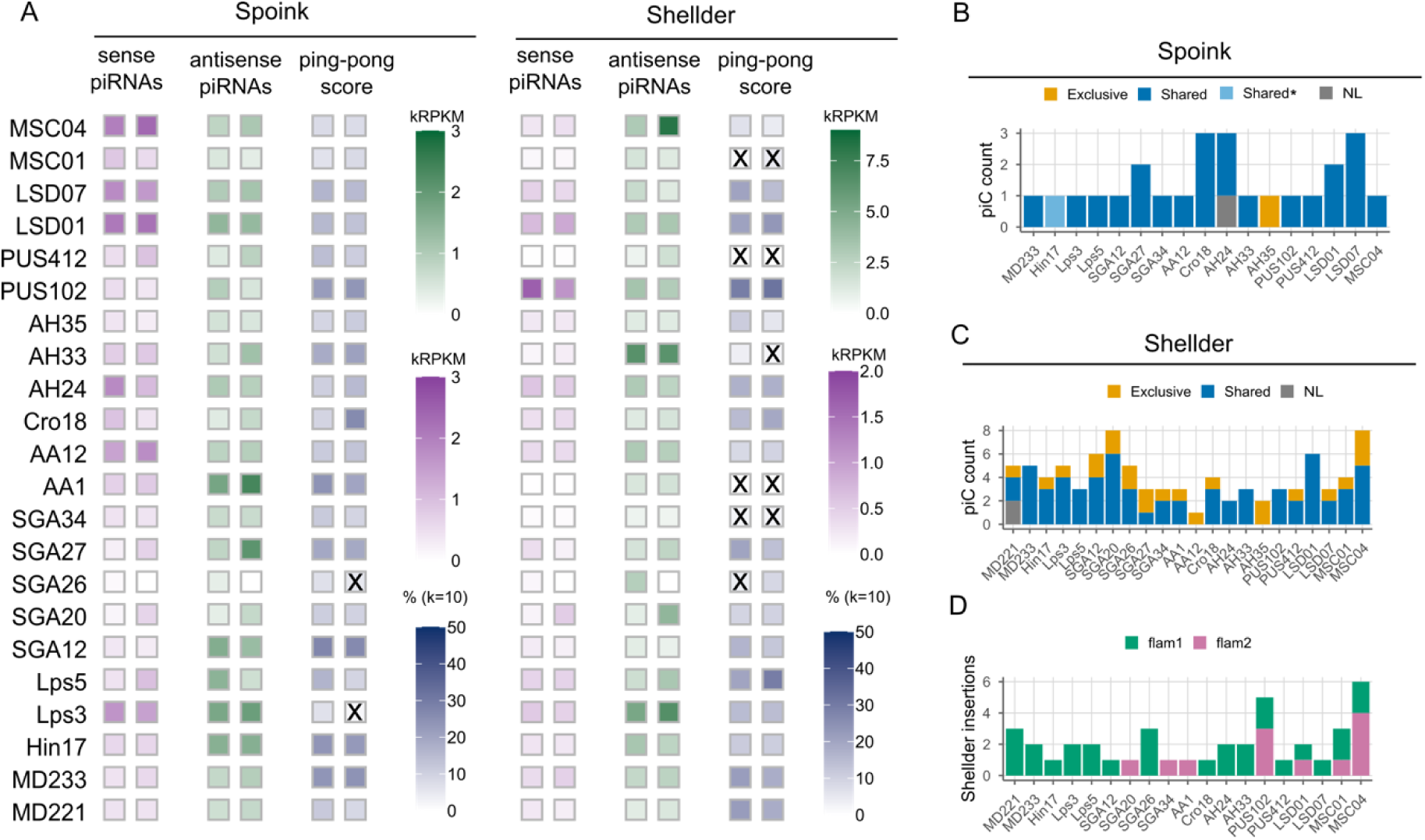
*Shellder* and *Spoink* defence. **(A)** Heatmaps of sense and antisense piRNA levels mapping to *Spoink* and *Shellder* are shown in kRPKM (thousand reads per kb per million mapped miRNA) and the ping-pong scores (fraction of 5′-to-5′ sense/antisense piRNA pairs that overlap by exactly 10 nucleotides out of all overlaps). Two biological replicates were plotted per strain, with random sampling when more than two libraries were available. Tiles marked with a cross (X) denote non-significant ping-pong signatures (pair count ≤ 10 or Z-score < 2) (**B, C**) Number of piRNA clusters harbouring *Spoink* and *Shellder* per strain, respectively, colored by status (Exclusive, Shared, Shared*, NL – non-lifted. Shared* -*Spoink* insertion in Hin17 is predicted as an exclusive cluster, but it is likely a shared cluster (see text). (**D**) Number of *Shellder* insertions found in *flam1* and *flam2* across the strains.

*Spoink* shows clear evidence of germline piRNA biogenesis, with all strains having both sense and antisense piRNAs, and evidence of a significant ping-pong signature (minimum pair counts = 10; Z-score >= 2; Fig. 4A; Fig. S14), consistent with previous work [65, 66]. For *Shellder*, a previous study found no ping-pong signature in the single strain examined [65]; across our larger sample, we detect a significant ping-pong signature in most, though not all, of our strains (18/22; minimum pair counts = 10; Z-score >= 2; Fig. 4A; Fig. S15). This result suggests that for most lines, the regulation of *Shellder* is not restricted to unistrand clusters expressed only in the soma, and that this TE may be active in the germline despite having an *env* gene.

### Shellder and Spoink insertions occur in established clusters

We also extended our test of the trap vs. *de novo* model to include *Shellder* and *Spoink*. The comparison between these LTR elements and the P-element is especially interesting because active LTRs may be more likely to form *de novo* piRNA clusters [59]. As the only strain uninvaded by these two TEs (the reference) was not included in our small RNA sequencing, we could not use the uninvaded strains to define established clusters as we did for the P-element. However, some clusters clearly pre-date these invasions: *flamenco*, for example, is an ancient cluster shared between *D. melanogaster* and *D. simulans* [60, 106]. Other clusters are shared among several strains from different geographic regions, which is only likely to be the case if their existence predates the invasion of these two TEs in the 1990s. We therefore considered these clusters as pre-existing, established clusters for this test. As for the P-element, we considered strains with insertions within annotated piRNA clusters as consistent with the trap model, and inconclusive otherwise.

For *Spoink*, 16 of the 22 strains had insertions (22 distinct, with three shared by two strains; n insertions = 25) in at least one cluster segregating at high frequency, and thus likely established as a piRNA cluster prior to the *Spoink* invasion (Figure 4B; Table S7). Note that one of these (Hin17), has an insertion in centromeric 500-bp satellite [116] with abundant cognate piRNAs in all of our strains (Fig. S16; cf.[117]; Given that the insertions lie in a TE-rich region, we thus considered this likely to be an established cluster despite being unable to assemble the region in other strains. For the remaining six strains, we were unable to find *Spoink* insertions in established piRNA clusters. In fact, we found annotated source loci for only one of them (AH35), which showed a production from an isolated, exclusive TE insertion; note that in this case, piRNA cluster annotation was supported by uniquely mapping reads, and not by the proTRAC annotation that may be susceptible to false positives. The remaining five strains may produce *Spoink* piRNAs from individual *Spoink* insertions consistent with *de novo* production, and/or clusters located in unassembled regions of the genome. For *Shellder*, we identified up to 95 independent insertions (107 in total) within piRNA source loci. Insertions in established piRNA clusters were found in 20 of the 22 strains (Figure 4C, Table S8), with many in *flamenco* (see below). In addition to these established clusters, several predicted piRNA source loci were exclusive and potentially *de novo* (23 of 58), including the only source loci identified in AH12 and AH35. However, most of the potentially de novo clusters for *Shellder* were detected only by proTRAC, and their contribution should be interpreted with caution. Overall, these results suggest that *D. simulans* has evolved to suppress both LTR elements through insertion into established piRNA clusters.

### Flamenco plays a major role in Shellder suppression

The primary source of piRNAs against retrovirus-like elements (LTRs with *env*) in *D. melanogaster* is *flamenco*, a large unistrand cluster on the proximal end of the X chromosome. Expressed in ovarian somatic follicle cells, *flamenco* predominantly produces piRNAs antisense to these elements, preventing them from mobilizing and invading the germline in this species [60, 108]. In *D. simulans*, the *flamenco* locus is duplicated (*flam1* and *flam2*) and enriched for LTRs that have recently invaded the genome [106]. Unlike *flamenco* in *D. melanogaster*, *flam1* and *flam2* do not appear to be restricted to somatic support cells, as they contain a large number of TEs predicted to be active in the germline, including many tandem R1 insertions and LTR elements that [106]. Moreover, the presence of uniquely mapping piRNAs in both sense and antisense orientations, together with detectable ping-pong signatures, suggests that *flam1* and *flam2* in *D. simulans* are dual-stranded and function in the germline as well as the soma [106]. In fact, no large solely unistrand clusters were found in *D. simulans* [114].

Here, we identified both *D. simulans flamenco* duplicates across all our strains, defined by their specific promoter sequences and downstream genes (see Supplementary Text). As expected, we find that *flamenco* in *D. simulans* appears to be very different from that of *D. melanogaster*. That is, while it does harbor retroviral-like elements, it also contains other types of TEs, particularly the non-LTR/LINE element R1 covering 89 ± 46 kb of *flam1* and 15 ± 12 kb of *flam2*. LTR insertions were predominantly antisense relative to *flamenco* transcription (83.6% in flam1 and 72.2% in flam2), whereas LINE insertions were predominantly sense (80.5% and 57.6%, respectively) (Fig. S17). We also find that most LTR families, including many with insertions in *flamenco*, produce ping-pong signatures (Fig. S18), suggesting germline expression of the element and, correspondingly, of the piRNA clusters that they inhabit. A few elements, however, do lack or have weaker ping-pong signatures in their cognate piRNAs, and either encode an *env* gene (e.g. NHMS-Beagle, Phanio) or belong to the MDG1 clade (e.g. Tabor, Stalker2), whose members encode a small transmembrane protein (sORF2) acting as a cell-cell fusogen that enables direct transfer of retrotransposon capsids from a somatic niche into the germline [118]. As expected, these elements show both a lack of ping-pong and a bias toward antisense piRNAs (Fig. S18). Note ping-pong signatures were computed for each TE family; we cannot determine whether this germline signal derives from *flamenco* or elsewhere in the genome.

Though the evolution of *flamenco* in *D. simulans* suggests it is no longer specialised for suppression of env-containing LTRs in somatic cells [106], we nevertheless find a high fraction of insertions of the *Shellder*, as did [65]: of the 95 putatively independent insertions, 40 were in the two duplicated *flamenco* loci, a significant enrichment (considering both *flam1* and *flam2* as 13.0% of the total length of piRNA cluster sequence; χ^2^ =71.004, df = 1, *p* < 2.2e-16; Fig. 4D). There was no apparent enrichment for *Spoink* insertions in *flamenco* (n-*flam* = 4, n-total = 26; χ^2^ = 0.129, *df* =1, *p* = 0.719). Due to the repetitive nature of the *flamenco* loci, however, we could not be sure that *Shellder* insertions in this region in different strains are truly independent using location alone. To accurately quantify the number of independent insertions, we therefore analyzed the flanking sequences, sense/antisense orientation, and SNPs shared between 5’ and 3’ LTRs (see Supplementary Text). This analysis shows that there are at least 28 independent insertions in *flam* (19 *flam1* and 9 in *flam2;* Table S9), constituting a significant enrichment (with both *flam* loci composing 13.0% of the weighted mean length of piRNA clusters (χ^2^ = 31.471, *df* = 1, *p* = 2.02e-08). In *flam1*, most of the independent *Shellder* insertions (11/19) occurred in R1 elements. These *Shellder* insertions are located mainly in two distinct microregions of R1, suggesting some target site specificity for this element [6 in 2483-2488 (↓CG↓CGCG) and 3 in 3943-3948 (↓CGTGCG)]. Interestingly, despite the presence of additional tandem R1 copies in *flam1*, there were no *Shellder* insertions in these locations. Such target site specificity is uncommon for LTR retrotransposons, although this has been recently reported for the invading *tirant,* which has been shown to integrate independently into the same nucleotide positions within host genes across different strains [56].

## Discussion

Here, we assembled genomes and annotated piRNA source loci for 22 *D. simulans* strains, both pre- and post-dating the P-element invasion of this species.

We show that P-element cognate piRNAs likely evolved primarily by the simple process of insertion of P-elements into existing piRNA clusters, as most post-invasion strains had P-element insertions in clusters already present before the P-element invasion. In the clear exception, piRNAs apparently originating from a P-element inserted antisense to a gene potentially producing double-stranded P-element RNA (dsRNA), is consistent with known mechanisms of *de novo* cluster formation [54, 56]. The earlier LTR invasions show a more mixed picture, with perhaps a larger role for *de novo* clusters. The reason may be that LTR retrotransposons, unlike the P-element, have a strong promoter, as well as a weak antisense promoter, and may yield some dsRNA on their own [119]. Nevertheless, for all three elements, clusters existing prior to their invasions seem to be a main source of cognate piRNAs, supporting the trap model.

Our test for the trap model, despite being one-sided, may be somewhat conservative as well. Expression of piRNAs can be epigenetically induced from cognate regions through generations [53]. Thus, the presence of an insertion in a piRNA cluster anywhere in the past history of a strain, or at low frequency within a strain, may have induced the piRNA production from individual TE insertions elsewhere in the genome.

Moreover, insertions into established clusters occurred repeatedly and mostly independently across strains, suggesting that resistance evolved largely in parallel across the population. This is perhaps unsurprising, as piRNA clusters comprise large regions of the genome, constituting a large target for transposition. Further, active, unsuppressed TEs will generate many new copies in a population per generation. In the case of the P-element in *D. simulans*, a single unsuppressed copy can transpose at high rates (0.0046 – 0.091/generation; [120]). A rough calculation shows that a single unsuppressed copy of a TE, transposing randomly at a rate of 0.5% per generation, has a 0.025% chance of inserting into a piRNA cluster, conservatively assuming that piRNA clusters constitute 5% of the genome. In *Drosophila*, with a large effective population size of around a million, this implies >2.5 new cluster insertions/generation at the point where the TE has spread to 1% of individuals. This rate is above the threshold needed to give rise to a soft selective sweep *via* multiple beneficial mutations [121, 122], in this case *via* multiple insertions into piRNA clusters [123]. Thus, the combination of a large target size for suppressive insertions, a large effective population size, and a high mutation rate is consistent with a regime of polygenic soft sweep adaptation (reviewed in [122, 124]). Consistent with this, even in laboratory populations, the P-element appears to be suppressed via insertion into piRNA clusters [27, 62], though in other cases it may be silenced *via* an emergent piRNA cluster [24, 63].

However, it may be that not all piRNA clusters confer equal protection against all TEs. Both piRNA clusters [125], and TEs [113] can show changes in expression throughout germline development, and their expression niches must overlap in order to result in TE silencing. We noted a preference for P-element insertions at the distal tip of the X chromosome (in telomeric associated sequences, X-TAS) and for the LTR element *Shellder* in *flamenco*. The telomeric P-element insertions are reminiscent of the Trans-Silencing Elements in *D. melanogaster,* in which a P-transgene inserted in into the tip of the X silences sequences elsewhere in the genome, later shown to involve piRNAs [126–129]. Moreover, naturally occurring P-element insertions accumulate in the piRNA cluster at the tip of the X chromosome in both natural populations and laboratory strains of *D. melanogaster* [60, 123, 126, 127, 130]. The reason may be that piRNA clusters, like some TEs, have particular expression niches [113]: the P-element is apparently active mainly during early germline development [131], and piRNA cluster homologous to the distal X-TAS cluster in *D. melanogaster* shows evidence of *trans-*silencing throughout the early stages of germ cell formation [53]. In *D. melanogaster*, the expression niche of *flamenco* is in the follicle cells, consistent with *env-*containing LTR elements infecting the ovarian germline from adjacent cells [108, 111, 113, 132, 133]. In *D. simulans*, *flamenco* appears to have additionally acquired germline expression, although large regions of *flam1* and *flam2* lack germline piRNAs [106]. *Flamenco* may therefore retain some follicle cell activity, as also suggested by the preponderance of recent *Shellder* insertions in these loci and by the antisense-biased piRNAs and the lack of ping-pong signatures of some <u>env</u>-containing and MDG1 clade elements. Our data suggest that *flamenco* retains an ancestral trapping role for retrovirus-like elements even as its structure and expression have diverged from those in *D. melanogaster* [106].

In summary, this work sheds light on the mode of evolution piRNAs cognate to newly invading TEs, likely underpinning adaptation to and silencing of these TEs. We may have missed some genuine piRNA source loci due to incomplete genome assembly or methodological limitations. We may also have missed transient, unstable piRNA source loci that might protect some strains early in a TE invasion—e.g., epigenetically induced silencing from individual TEs, triggered by maternal inheritance of piRNAs [53]. Nevertheless, the consistent recruitment of pre-existing clusters to piRNA production across three unrelated invading TEs suggests that this is an important mode of adaptation to TE invasions, allowing hosts to respond rapidly to sequences that they have never encountered.

## Acknowledgements.

We thank G Lee and D Presgraves for helpful comments on an earlier version of this manuscript, and K Senti for discussion. The piRNA data generation was carried out by the Centre for Genomic Research, which is based at the University of Liverpool. We acknowledge the support of Dr. Richard Gregory and the Centre for Genomic Research (CGR) at the University of Liverpool for providing access to computational resources used in this research.

## Supplementary Information

### Supplementary Materials and Methods

#### High molecular weight (HMW) DNA isolation

We isolated HMW DNA using either a standard 25:24:1 phenol-chloroform-isoamyl (PCI) protocol or a Qiagen Genomic Tips 20/G HMW kit (Qiagen) following the manufacturer’s protocol. For PCI extractions, 10 females/line were homogenized in a glass Dounce homogeniser (Sigma-Aldrich) with liquid nitrogen, and incubated in 0.7 ml of cell lysis buffer with proteinase K (Qiagen) in 1.5 ml LoBind Eppendorf tubes at 50°C for 2 hours. We added 0.75 ml of PCI, inverted several times, and then spun them (∼14000 rpm) for 20 mins at 4°C. The upper phase (0.5 ml) was transferred to a fresh 1.5 ml tube with 0.5 ml of chloroform, spun again (∼14000 rpm) for 20 mins at 4°C, and transferred to a new tube with 0.5 ml of cold isopropanol (-20°C) and 35.7 μl 3M NaOAc added. The samples were inverted several times, placed on ice for 20 mins, and then spun (∼14000 rpm) for 30 mins at 4°C. Finally, the samples were washed with 1 ml of cold 80% EtOH (-20°C), inverted and spun (∼14000 rpm) for 5 mins at 4°C. The DNA pellets were air dried and resuspended in 0.1 ml of nuclease-free H2O.

From 5-10 extractions from the same line were were pooled, RNAse A treated and cleaned by concentrated to a volume of 0.2 ml, adding 20 μl of RNAase A (Thermo Fisher Scientific) and incubating at 37°C for 30 mins. Samples were re-precipitated with 22 μl of 3M NaOH and 0.55 ml of 100% EtOH, placed at -80°C for 30 mins and then spun (∼14000 rpm) for 30 mins at 4°C. The samples were then washed with 80% EtOH (-20°C), inverted and spun (∼14000 rpm) for 5 mins at 4°C before being air dried and resuspended in 0.4 ml of nuclease-free H2O. We eliminated short fragments from our DNA isolates by doing a 0.8X clean-up with AMPure XP beads (Beckman Coulter). The samples were incubated with beads at room temperature (RT) for 10 mins and pelleted using a magnet. The eluate was removed and discarded and the pellet washed twice with 70% EtOH before being air dried. The pellet was resuspended in 0.1 ml of nuclease-free H2O and incubated at 37°C for 10 mins. The beads were placed back on the magnet and then eluate removed and placed in a new 1.5 ml tube. The DNA concentration was measured with Qubit (Thermo Fisher Scientific), and DNA integrity checked on a Bioanalyzer (Agilent).

#### Genomic Tips

We isolated DNA from pools of 20 female flies per line using the Genomic Tips 20/G HMW kit (Qiagen), following the manufacturer’s protocol. Briefly, the flies were ground to a fine powder using a pestle and mortar and liquid nitrogen. The ground flies were transferred to pre-warmed lysis buffer with 20 mg/ml proteinase K and 10 mg/ml RNase A and incubated for a further 2 hours at 50°C. The samples were spun (∼7400 rpm) for 10 mins at 4°C and the supernatants were transferred to new tubes, vortexed for 10 s and then promptly loaded onto the columns, whereby they drained by passive gravity until empty. The columns were washed twice with 1 ml of QL buffer, and the DNA eluted into 1.5 ml Eppendorf LoBind tubes with 1 ml of warm QF buffer. Each sample was precipitated with 0.7 ml of isopropanol and mixed via tube inversion. Each elution was then spun (∼12700 rpm) for 30 mins at 4°C. The resulting pellets were washed with freshly prepared 70% EtOH and spun (∼14000 rpm) for 10 mins at 4°C. After air drying, the DNA pellets were resuspended overnight at 4°C in 50 μl of nuclease-free water. Libraries for Nanopore sequencing were prepared the following day.

#### Library preparation and Nanopore sequencing

*SQK-LSK109 kit*. Libraries for long read sequencing were prepared using the Ligation Sequencing kit, SQK-LSK109 (Oxford Nanopore), according to the manufacturer’s protocol. Approximately 2.5 μg of DNA was used per library preparation. The total volume of the DNA was adjusted to 49 μl with nuclease-free water. We performed a DNA Repair and end-prep step, where DNA was mixed with NEBNext FFPE DNA Repair Buffer, NEBNext FFPE DNA Repair Mix, Ultra II End-prep reaction buffer and Ultra II End-prep enzyme mix and incubated in a thermocycler for 5 mins at 20°C followed by 5 mins at 65°C.

The DNA was cleaned and size-selected using AMPure XP beads (Beckman Coulter). The DNA underwent adaptor ligation, where ligation buffer, NEBNext Quick T4 Ligase and Adapter Mix was added and the sample incubated for 10 mins at RT. The samples were cleaned and size-selected again using AMPure XP beads and Long Fragment Buffer and resuspended in Elution Buffer. The libraries were mixed with Sequencing Buffer and Loading Beads before being loaded into prepared R9.4.1 flow cells which had, on average, 1000 active pores. The flow cells were run for 72 hours or until there were no pores available, in a MinION.

*SQK-LSK114 kit.* Libraries from the Ligation Sequencing kit, SQK-LSK114 (Oxford Nanopore) were prepared in much the same way as above, according to the manufacturer’s protocol. Approximately 1.5-2 μg of DNA was used per library preparation. The total volume of DNA was adjusted to 47 μl with nuclease-free water and the DNA Repair and end-prep steps were performed. The DNA was cleaned and size-selected with AMPure XP beads (Beckman Coulter) and then underwent adaptor ligation before being cleaned and size-selected again. Libraries were loaded into prepared R10.4.1 flow cells and run on a GridION (Centre of Genomic Research, Liverpool) for 72 hours.

#### RNA extraction, library prep and smallRNA sequencing

We performed small RNA sequencing on all 22 lines (TableS1, *n* = 2-6 replicates). For each library, we dissected ovaries from 3-5-day-old females frozen at -20°C in RNA*later*-ICE (Ambion), and extracted total RNA as follows: dissected ovaries were pooled in groups of 10 per line and immediately homogenized in 0.5 ml TRIzol Reagent (Invitrogen) using a tissue homogenizer (bertin technologies) and incubated on ice for 5 mins. We added 0.2 ml chloroform (Sigma) and inverted each sample for 15 s before spinning them (12000 rpm) for 15 mins at 4°C. The top aqueous phase was moved into a new tube with 0.5 ml isopropanol. The samples were incubated at RT for 10 mins and spun (12000 rpm) for 10 mins at 4°C. Finally, the samples were washed in 1 ml 75% EtOH, air-dried and resuspended in 50 μl RNAse-free water.

RNA concentration was measured with Qubit (Thermo Fisher Scientific), and RNA integrity checked on a Bioanalyzer (Agilent). Prior to small RNA-seq library prep (CGR, Liverpool), a DNAse treatment was performed on each sample. Samples were then cleaned and concentrated using a column-based kit (Zymo). Libraries were prepared using the NEBNext small RNA library prep kit, including an additional 2S rRNA blocking step (as described in Wickersheim and Blumenstiel 2013). The libraries were sequenced on one lane of the Illumina NovaSeq X Plus using 10B chemistry at the Centre for Genomics Research in Liverpool.

#### Genome assembly and scaffolding

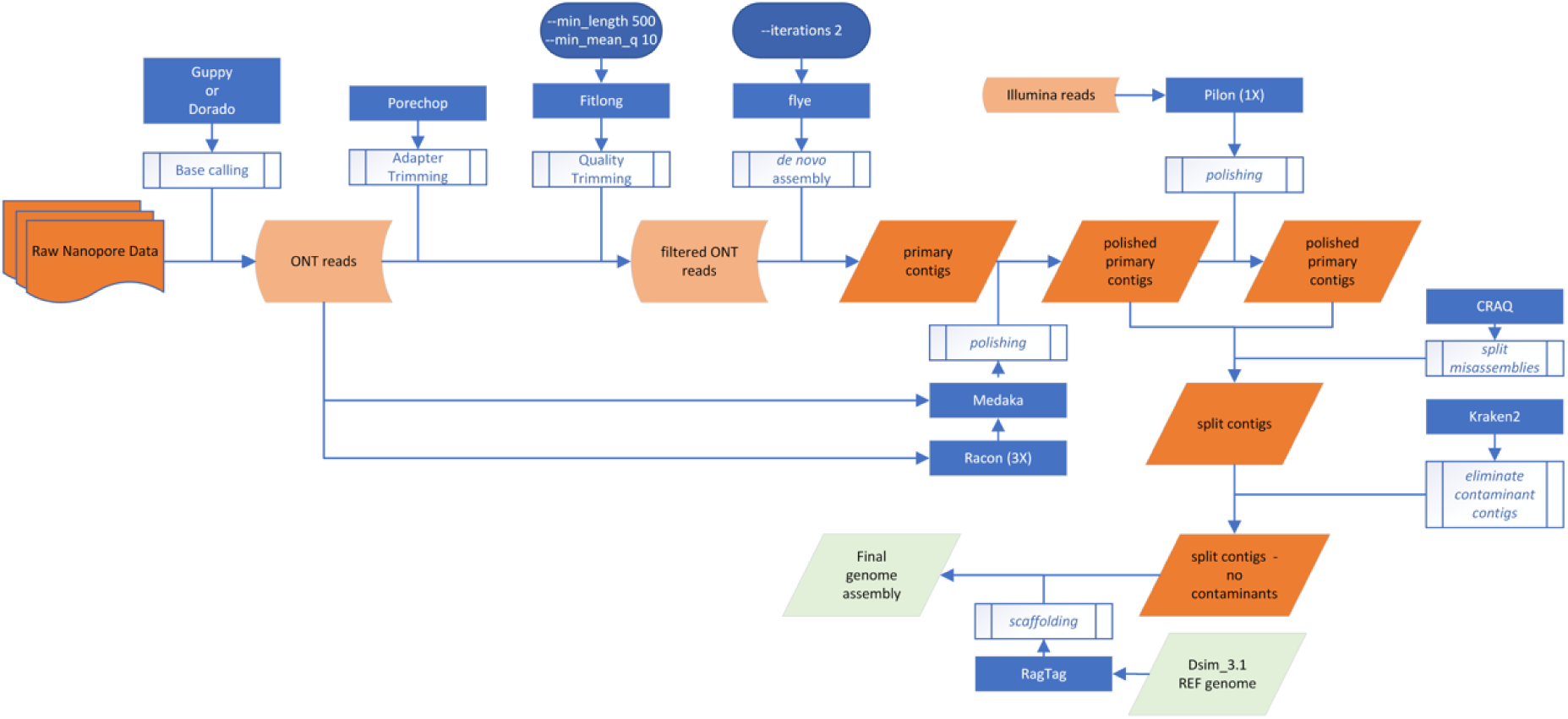

Nanopore reads were base called using Guppy v3.2.10 (Oxford Nanopore Technologies PLC) or Dorado v0.7.2 (Oxford Nanopore Technologies PLC). Porechop v0.2.4 (https://github.com/rrwick/Porechop) was used to trim adapters and Filtlong v0.2.1 (https://github.com/rrwick/Filtlong) for quality filtering (--min_length 500 --min_mean_q 10), and NanoPlot v1.42.0 [134] to obtain summary statistics describing the reads. An initial *de novo* assembly was obtained using Flye v2.9.2 [69] with two iterations of polishing with the raw reads. Additional polishing was performed using raw nanopore reads with three rounds of Racon v1.5.0 [70] and Medaka v1.11.3 (https://github.com/nanoporetech/medaka). For Lps5, Hin17, Cro18 and SGA27, Illumina reads were available, and Pilon v1.24 [71] was used to polish these genomes with Illumina reads (1 round). To detect potential misassemblies, we used Craq v1.0.9-0 [72] to detect structural errors and break chimeric contigs. Kraken2 v2.1.3 [73] was then used to identify and eliminate contigs from potential contaminant sources.

To scaffold these *de novo* assemblies on a reference genome, we used RagTag v2.1.0 and the Dsim_3.1 (GCF_016746395.2) reference genome assembly. Several tools were used to assess the quality of genome assemblies: 1) BUSCO v5.7.1 [75] was used to assess the gene content using the diptera_odb10 database. 2) Quast v5.2.0 [76] was used to compute basic assembly statistics, such as total length, contig numbers, and N50. 3) Craq v1.0.9-0 was used to quantify clip-based regional and structural errors (Table S3).

#### D. simulans TE library curation

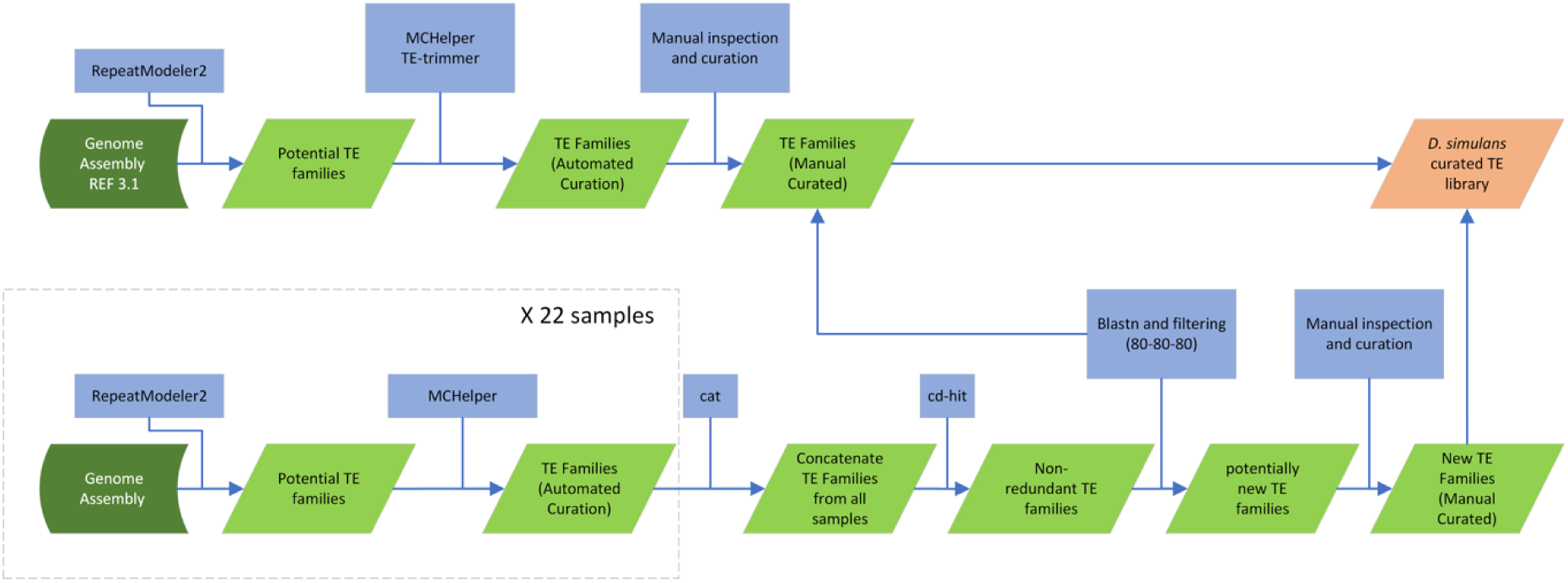

RepeatModeler v2.0.3 [79] was used to identify potential TEs and other repeats in the reference *D. simulans* genome Prin_Dsim_3.1 (GCF_016746395.2). We used MCHelper v1.7.0 (https://github.com/GonzalezLab/MCHelper) with default parameters and fully automated analysis, or TE-trimmer v1.4.0 (https://github.com/qjiangzhao/TEtrimmer) with default parameters to curate consensus sequences. TE-aid (https://github.com/clemgoub/TE-Aid) was used to inspect each consensus sequence, and problematic sequences were further curated manually, according to the TE manual curation guide [84]. In some cases, an online BLAST self-alignment (using the same sequence as both query and subject, with the parameters “align two or more sequences” and “somewhat similar sequences”) was employed to help determine the boundaries of the TE families, either by identifying their terminal repeats or target site duplications (TSDs).

We then obtained additional TE sequences from the 22 *D. simulans*. RepeatModeler was run for each additional genome, followed by MCHelper. The MCHelper consensus sequences of each sample were concatenated into a single multifasta file, and the redundancy was eliminated using cd-hit (-d 0 -aS 0.8 -c 0.8 -G 0 -g 1 -b 500). The remaining sequences were then blasted against the curated library from the reference (-outfmt "6 qseqid sseqid pident length mismatch qstart qend qlen slen" -evalue 1e-20 -reward 2 -penalty -3 -gapopen 5 -gapextend 2 - max_target_seqs 1), to eliminate redundant, already curated sequences. Any sequences that do not meet the criteria of 80% identity over 80% query sequence coverage were inspected and curated as described above.

Finally, we ran pantera, which identifies polymorphic TEs from pangenome data [82], to identify TE sequences contributing to insertion/deletion polymorphism. The same blast and curation strategies described above were used to eliminate already curated TEs and curate potential new sequences, with four sequences identified in this way. The final curated *D. simulans* library consists of 224 sequences.

#### Combined Drosophila TE library curation

A combined *Drosophila* TE library was obtained from: 1) 224 sequences from the *D. simulans* library described above; 3) 157 sequences from a modified version of the MCTE library [85], supplemented with 5 additional sequences from Repbase23.11 [86] and the BDGP TE library [30], 3) 2,263 sequences from diverse *Drosophila* species [87], inspected to eliminate satellites, rDNA, Low-Complexity sequences, and clear chimaeras or non-TEs. The final combined TE library comprises 2,649 sequences.

#### TE annotation of genome assemblies

We identified the locations of TE sequences in our genome assemblies using two main modules. In the first, we use the curated *D. simulans* TE library to annotate TE families with high sequence similarity. In the second, we employ the combined *Drosophila* TE library to annotate divergent TE fragments.

**Figure 1:**
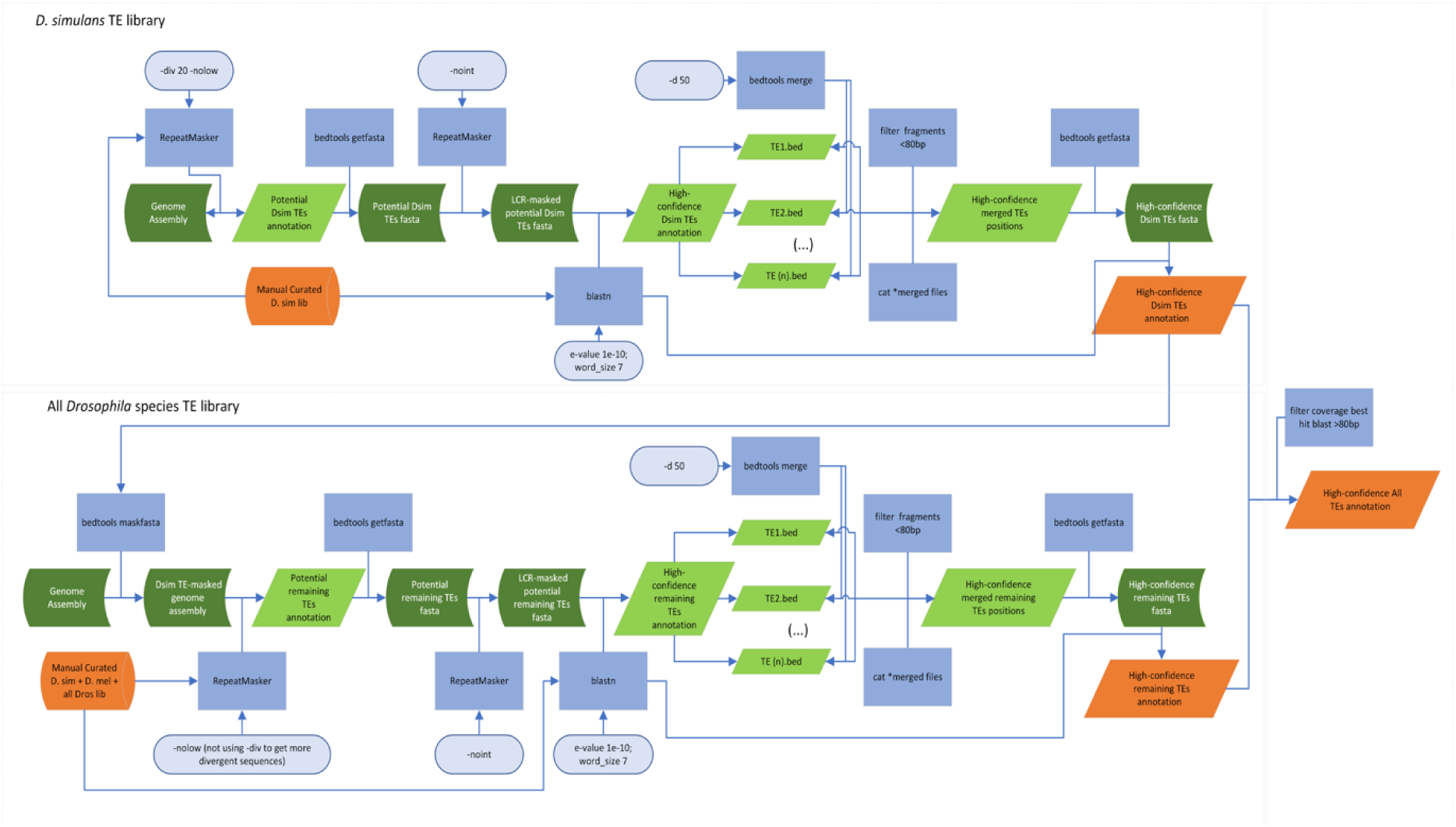
Workflow of TE annotation. The steps in the workflow are described in detail in the main text.

Module 1: Genome assemblies were initially annotated using RepeatMasker [88] with the curated *D. simulans* TE library, employing -div 20 to recover conserved sequences and -nolow to avoid fragmentation of complete copies. Potential TE fragment sequences were retrieved from the respective genomes using getfasta from bedtools v2.30.0 [78] based on the RepeatMasker annotation. To retain only reliable TE fragments, they were further masked for simple repeats and low-complexity regions using RepeatMasker (-noint) and a blastn search was performed against the curated *D. simulans* TE library (-e-value 1e^-10^; -word-size 7; - max_target_seqs 1; -max_hsps 1), eliminating sequences with no significant matches. Finally, potential split fragments were merged using bedtools merge with a window size of 50 (-d 50) and fragments less than 80 bp were excluded. An additional step of blastn is performed against the curated library to retrieve the complete TE classification, generating a high-confidence annotation of *D. simulans* more conserved TEs.

Module 2: The annotated TEs from Module 1 were masked in the genome assemblies using bedtools maskfasta. The masked genome was submitted to RepeatMasker (-nolow) to annotate the remaining more divergent TE fragments using the combined *Drosophila* TE library. The same steps from Module 1 were performed to obtain a high-confidence annotation of the remaining TE fragments. Finally, the annotations from the two modules were combined.

To estimate the number of TE copies, we used the set of scripts “One code to find them all” [90] to parse the RepeatMasker results obtained with the curated *D. simulans* TE library.

#### Reads filtering

Short reads were trimmed with Cutadapt v1.2.1 [91] to remove adapters and select reads of 23-29 nucleotides. Reads mapping to annotated rRNA, tRNAs, miRNA and miscRNAs were eliminated using Bowtie v1.3.1 (-v 2 -k1 -y -un)[92]. The different sets of RNAs were downloaded from Flybase for *D. melanogaster*. The remaining reads were then mapped to *D. simulans* mRNA sequences (GCF_016746395.2_Prin_Dsim_3.1_rna.fna) with Bowtie v1.3.1 (-n 1 -l 12 -a --best --strata -y). Reads mapping the sense strand of genes were removed to eliminate potential small RNAs derived from degraded RNAs. Because many TEs are located within introns, small RNAs mapping to TEs (using BWA aln) were added back to the dataset. TBr2_collapse and TBr2_duster from the NGS toolbox [93] were used to collapse the remaining reads to include only unique piRNAs and to remove low-complexity reads.

#### piRNA cluster prediction - Protrac protocol

Filtered reads from each sample were mapped to their respective genome assemblies, allowing multi-mapping with Bowtie v1.3.1 (bowtie -n 1 -l 12 -a --best --strata -y --chunkmbs 1024 -S). Next, proTRAC_2.4.3 [93] was run on sam files (-swsize 500 -swinger 100 -size 500 -1Tor10A 0.6 -min 23 -chitin 10 -pens 0.05 -pti).

#### piRNA cluster prediction - Coverage-based protocol

Filtered reads from each sample were mapped to their respective genome assemblies, allowing only unique mapping. To maximise the number of unique mappings, we performed two rounds of mapping using Bowtie v1.3.1. In the first round, no mismatches were allowed (bowtie -n 0 -l 12 -a -m 1). In the second round, the unmapped reads from the first round (recovered with samtools v1.15.1 fastq function) were remapped with one mismatch (bowtie -n 1 -l 12 -a -m 1). Finally, we used samtools merge to combine the SAM files. Windows of 500 bp were created with bamCoverage v3.5.5 of deepTools [135] to calculate normalised counts (CPM). Windows with an expression of at least 8 CPM were filtered and merged if they were located within 10 kb of each other using samtools. RPKM values were then calculated for the resulting regions, and those with an RPKM value of at least 25 were selected. To analyse the 1U bias, the genomic regions were extracted using bedtools getfasta, and the reads were remapped to these regions with the same parameters. The BAM files were subsequently divided into separate files and we used FastQC v0.12.1 (https://github.com/s-andrews/FastQC) to produce a quality control report for each of them from where the 1U proportion of each region was extracted. Clusters with 1U frequencies above 60% were maintained.

#### Merging and filtering piC annotations within strains

For each strain, piRNA clusters predicted from different replicates were first merged using a 10kb window, with merging performed separately for each of the two approaches above. Then, predicted clusters from both approaches were combined and merged for each strain using bedtools merge. Clusters intersecting with genes were split to eliminate gene regions using bedtools subtract. The gene annotation for each strain was obtained using Liftoff v1.6.3 [136], transferring the current gene annotation of the Prin_Dsim_3.1 genome (GCF_016746395.2). AGAT v1.4.1 (https://github.com/NBISweden/AGAT) was used to filter the reference GFF file prior to liftoff, removing genes corresponding to lncRNAs (detected through string search) and TEs annotated as genes (identified via blastn of the curated TE library against exons). Clusters that did not intersect with the TE or were less than 2kb were removed.

#### Converting coordinates and checking shared piCs

For each strain, piC coordinates were transferred to the reference genome (Prin_Dsim_3.1) using the halLiftover tool [94] with the hierarchical alignment (HAL) obtained by the pangenome analysis. Further efforts to acquire additional transferred coordinates included using halLiftover for 10 kb upstream and downstream piC regions, as well as Liftoff. All lifted coordinates across strains were then intersected with bedtools merge to identify shared piCs.

#### Ping-pong signal

Filtered reads (size-selected and depleted of annotated non-coding RNAs, as described above) were mapped to the curated *D. simulans* TE library or to the *Tctp* reference sequence (GD18736) using Bowtie v1.3.1 (https://github.com/BenLangmead/bowtie), allowing up to 2 mismatches. To quantify the ping-pong signature, we used the Python script signature.py (https://github.com/ARTbio/tools-artbio/blob/main/tools/small_rna_signatures/signature.py), using the method originally described by [95]. A ping-pong signal was considered significant when the 10-nucleotide overlap had at least 10 supporting read pairs and a Z-score ≥ 2.

#### Detection of unassembled TE copies from long reads

Filtered long reads from the 22 strains were mapped to the reference genome (GCF_016746395.2) using minimap2 [97] and Sniffles2 [98] was used to call structural variants from the alignments. Custom Python and Bash scripts were used to extract insertion sequences, identify *P, Shellder,* and *Spoink* insertions via blastn, and extract the TSD sequences associated with each insertion.

### Flamenco genes delimitation in *D. simulans* strains

All *D. simulans* strains sequenced here have two copies of *flamenco* (*flam1* and *flam2*), as expected from previous work [137]. In the reference genome, *flam1* is found on chromosome X (NC_052525.2), while *flam2* is located in an unplaced contig (NW_025416856.1).

We defined the boundaries of *flam1* (NC_052525.2:21,072,908-21,518,427) from the promoter region to the first downstream *Dip1* gene. In our pipeline, *flam1* was recognised as two piRNA clusters (813, 814). The split of *flam1* into two clusters is due to an interruption by a gene in the reference genome and in several other strains. This gene corresponds to the *translationally-controlled tumor protein homolog* (*Tctp* - LOC120285394), a highly conserved eukaryotic gene that plays crucial roles in organ growth, cell survival, and DNA repair, among others [138, 139]. Analysis of the *Tctp* copy in the reference genome reveals a poly (A) tail in the 3’ region, suggesting the copy was inserted by retroposition of its mRNA. While *Tctp* is a single-copy essential gene in *D. melanogaster*, our sequence analysis revealed that it has been duplicated in the vast majority of the *D. simulans* strains analysed.

The boundaries of *flam2* in the 3’ regions are harder to establish. We defined the gene from the promoter sequence until LOC6740432 (antigen 5-like allergen Cul n 1) (NW_025416856.1:19298-442148). It encompasses the clusters 870, 871 and 872 identified in our pipeline. It is interrupted by two uncharacterized protein genes (LOC6740449 and LOC27208153). Both genes are homologous to the *pirate* gene (*pira*), which encodes a SUMO protease. In *D. melanogaster*, the pirate gene is located on the X chromosome (X:12,760,642-12,763,611) and is silenced in male testes by piRNAs produced from a specific Y chromosome locus called *petrel* [140].

#### Independence of Shellder insertions in flamenco

For the P-element, independent insertions are relatively easy to identify as each insertion generates an 8-bp TSD. For *Spoink* and *Shellder,* whose TSDs are shorter (4-bp), this distinction is less straightforward. *Spoink* insertions nevertheless carried diverse TSDs and could be readily distinguished. In contrast, most *Shellder* insertions had the same TSD (CGCG; 40 of 69 insertions), so additional features were needed mainly due to the high insertion density in flamenco and insertions in the same repetitive TE (e.g. R1copies). We quantified the number of independent *Shellder* if *(i)* they contained different flanking sequence immediately adjacent to the insertion (including the target-site duplication), as these are generated from the target DNA upon insertion, or *(ii)* they are in reverse orientation from other insertions in the same region/target sequence, or (*iii)* they contain different alleles at a SNP shared between their own 5’ and 3’ LTRs. Because the two LTRs of an LTR retrotransposon are identical upon insertion, a SNP present in both LTRs of a copy must have been present at the time of insertion and different alleles at such SNPs therefore indicate distinct insertion events. Insertions that could not be distinguished by any of these criteria were conservatively considered a single insertion, so that our estimate represents a minimum number of independent insertions (Table S9).

---

**Figure S2:**
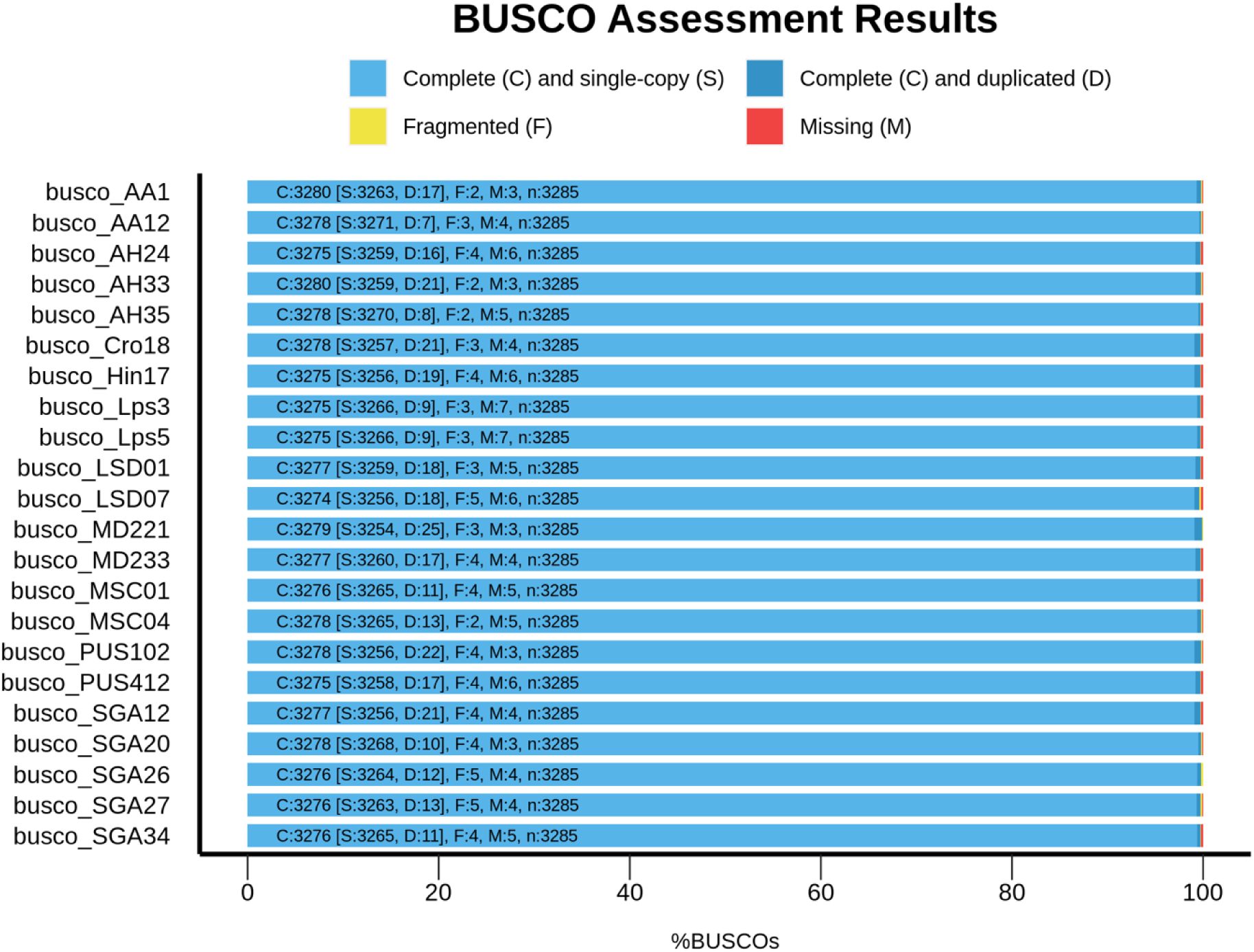
BUSCO completeness profiles for each genome, showing the proportion and number of complete single-copy (S), complete duplicated (D), fragmented (F), and missing (M) orthologs. Values are based on a total of 3,285 conserved genes.

**Figure S3:**
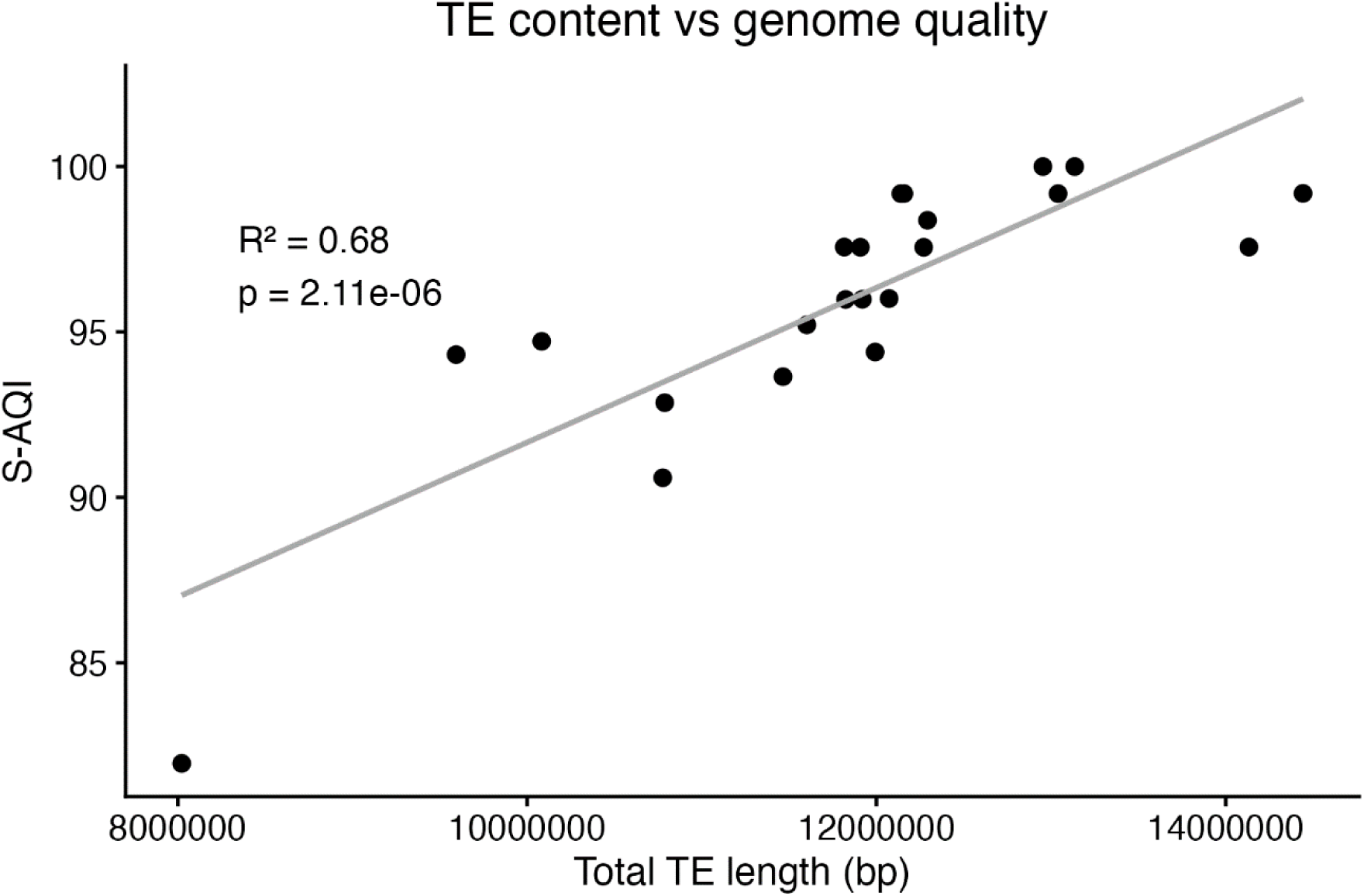
Correlation between total TE length (bp) and genome quality (S-AQI) for all. S-AQI quantifies genome assembly quality based on Clip-based Structural Errors detected by CRAQ (Li et al. 2023).

**Figure S3:**
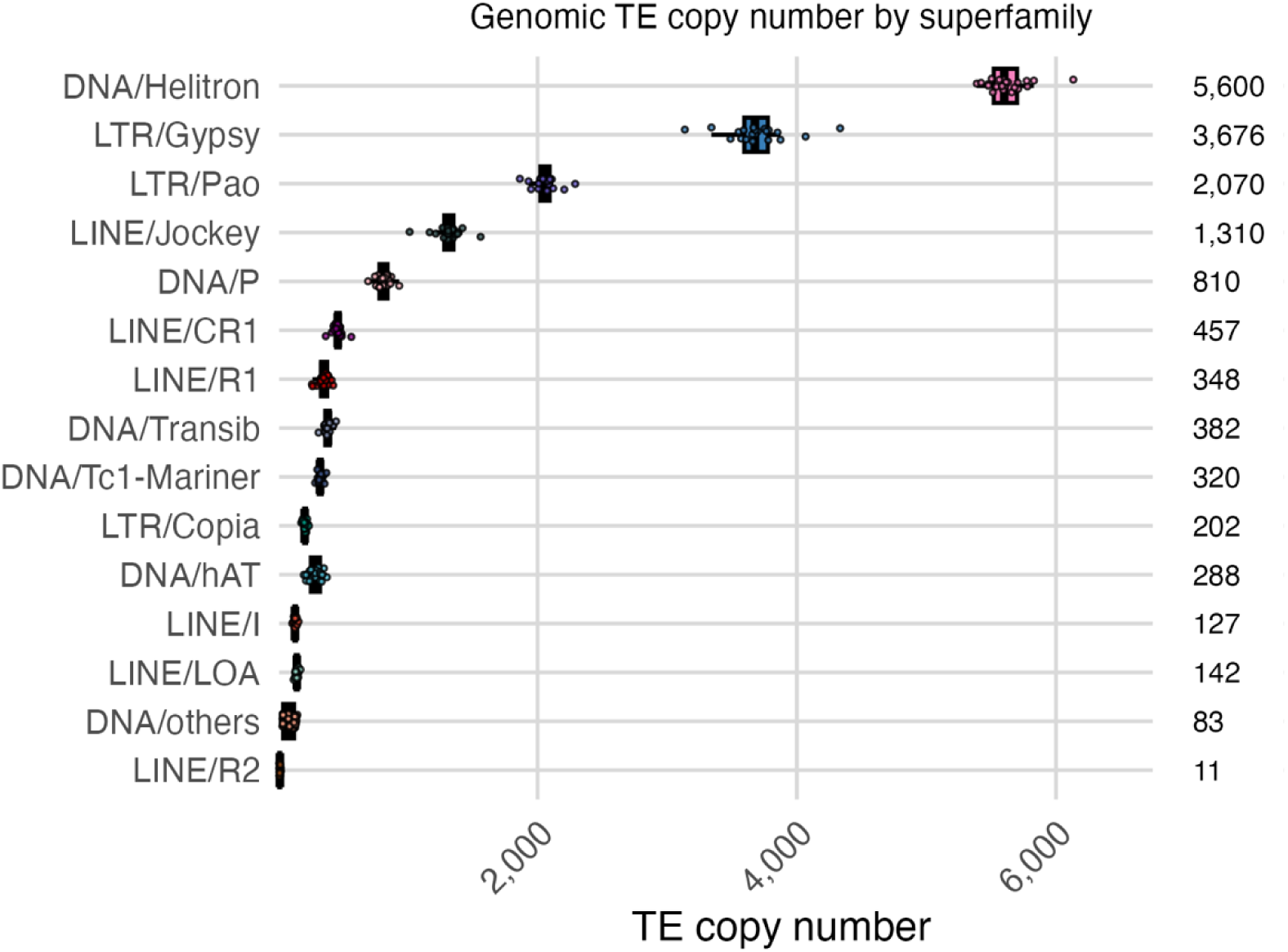
Distribution of copy numbers across the 22 D. simulans strains and the reference genome for each TE superfamily, as estimated by the One Code to Find Them All pipeline. Median values are shown on the right. Note that the ranking here differs from that of Mérel et al., 2020, due to a ∼5-fold smaller number of annotated Jockey copies. This difference is perhaps due to the refinement of the Jockey consensus sequence in our analysis, which involved removing repetitive sequences that likely belong to the insertion site in the genome rather than the element itself.

**Figure S4:**
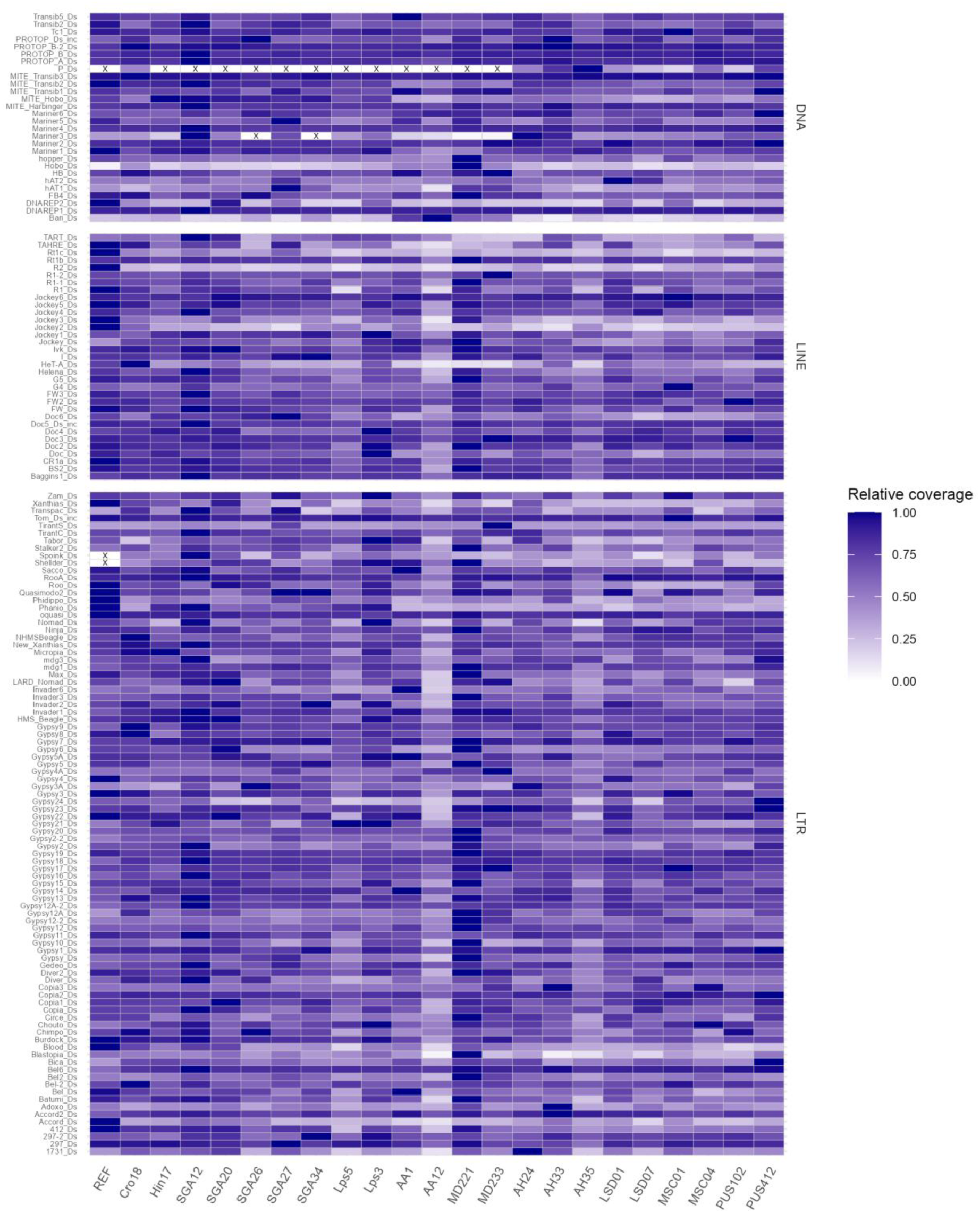
Relative coverage of TE families across strains. Values are normalised per family (maximum *= 1.0). “X” indicates complete absence. Consensus sequences that represent TE fragments are not shown. DNAREP1 subfamilies were summed into a single family*.

**Figure S5:**
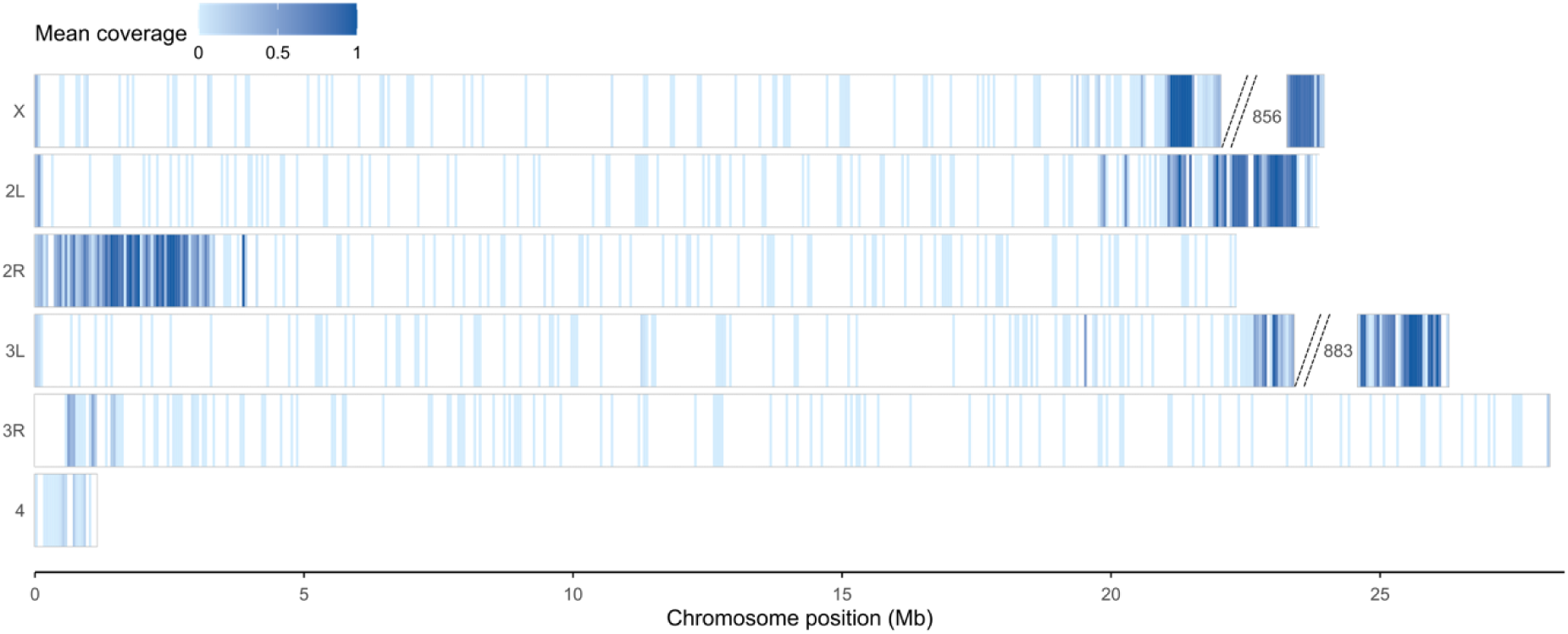
Genome-wide distribution of piRNA cluster coverage across chromosomes. Heatmaps represent mean piRNA cluster coverage across strains along the reference chromosomes, calculated in 50-kb windows. 856 corresponds to the contig NW_025416856.1, which belongs to chromosome X and contains the flam2 gene. 853 corresponds to the contig NW_025416883.1, which probably belongs to the end of chromosome 3L, as it contains genes orthologous to D. melanogaster genes located at the end of 3L.

**Figure S6:**
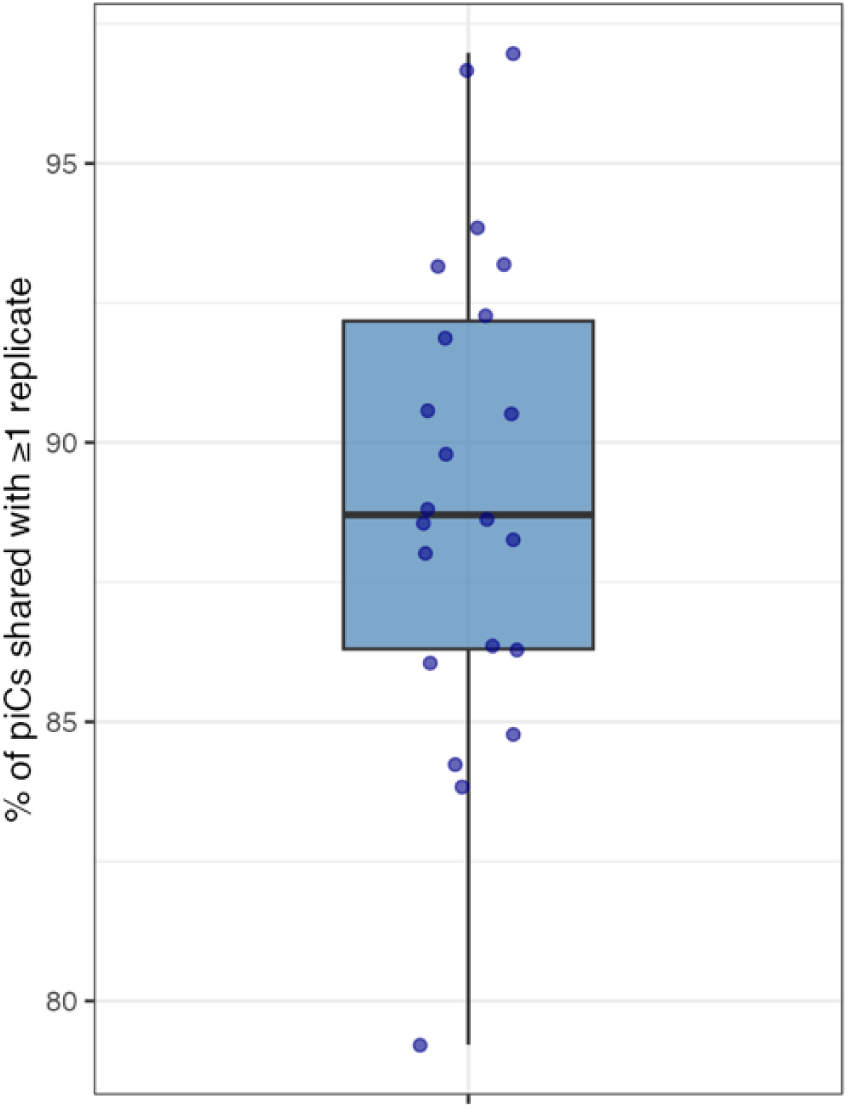
Percentage of piRNA clusters predicted that are shared across replicates per strain.

**Figure S7:**
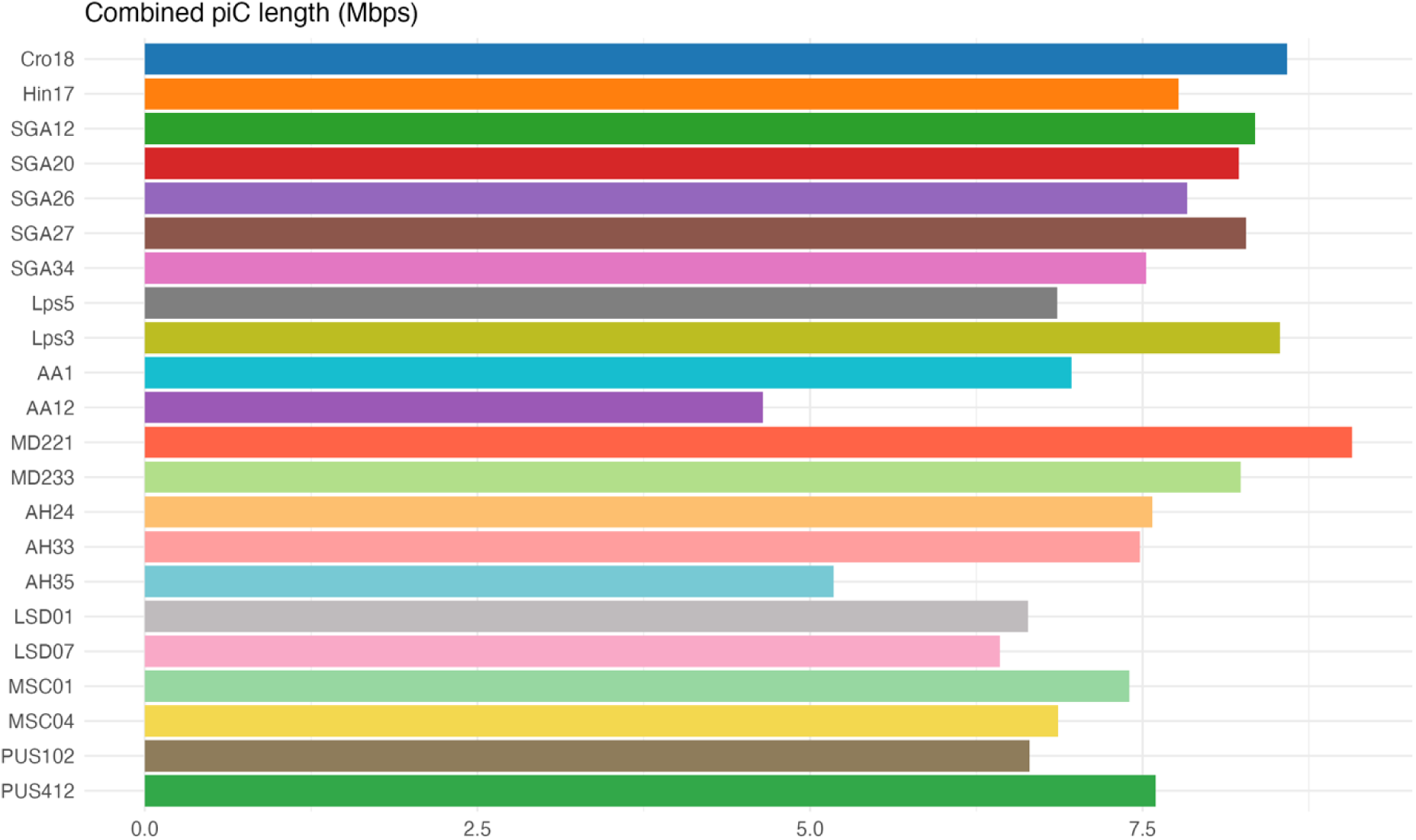
Combined piRNA cluster lengths per strain (Mb), averaged across replicates.

**Figure S8:**
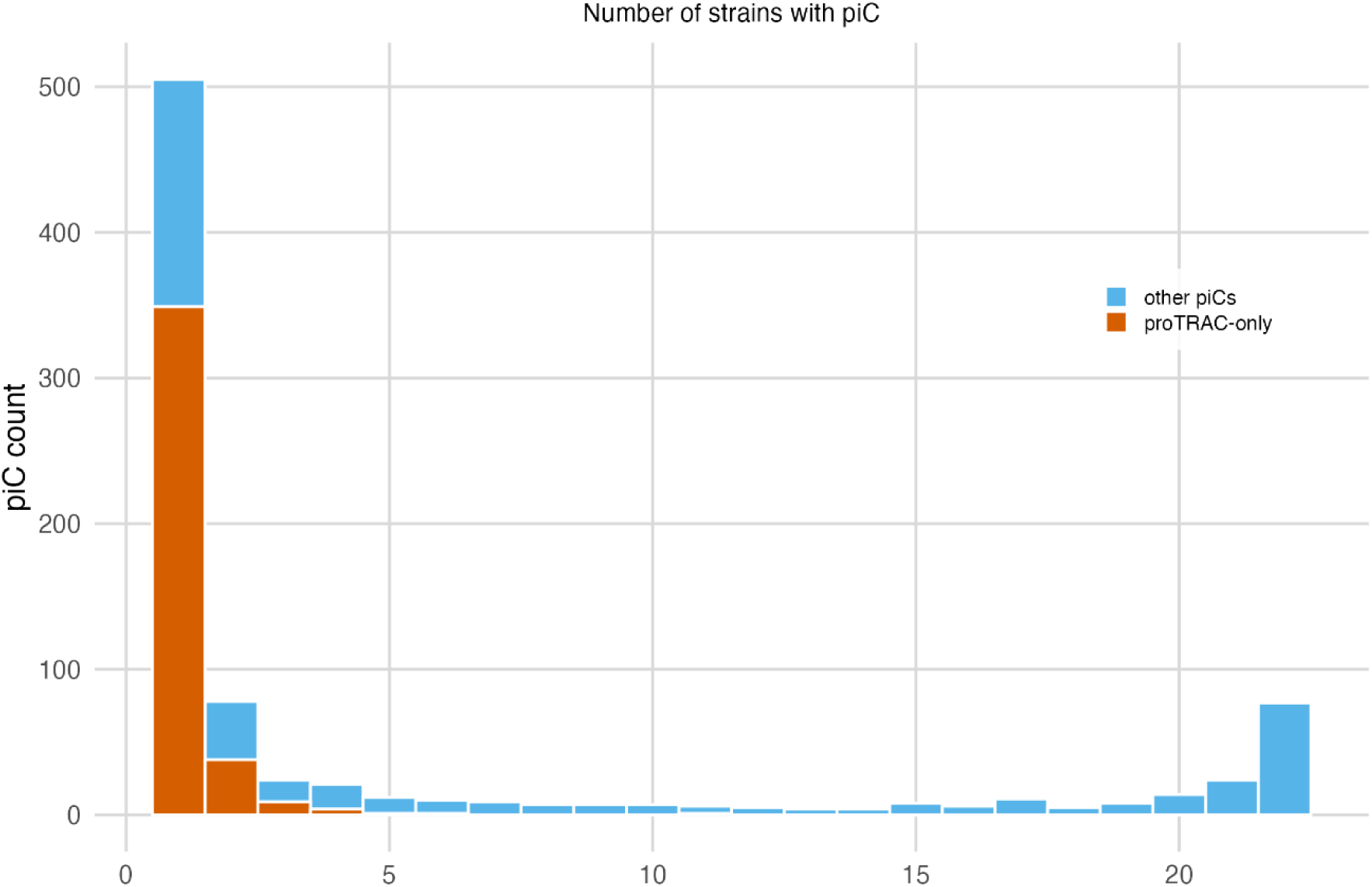
Distribution of all piRNA clusters (852) according to the number of strains in which they are detected, including those in regions that were not assembled in all strains. Clusters predicted by proTRAC-only are shown in a different colour from the others (predicted by coverage-only or both).

**Figure S9:**
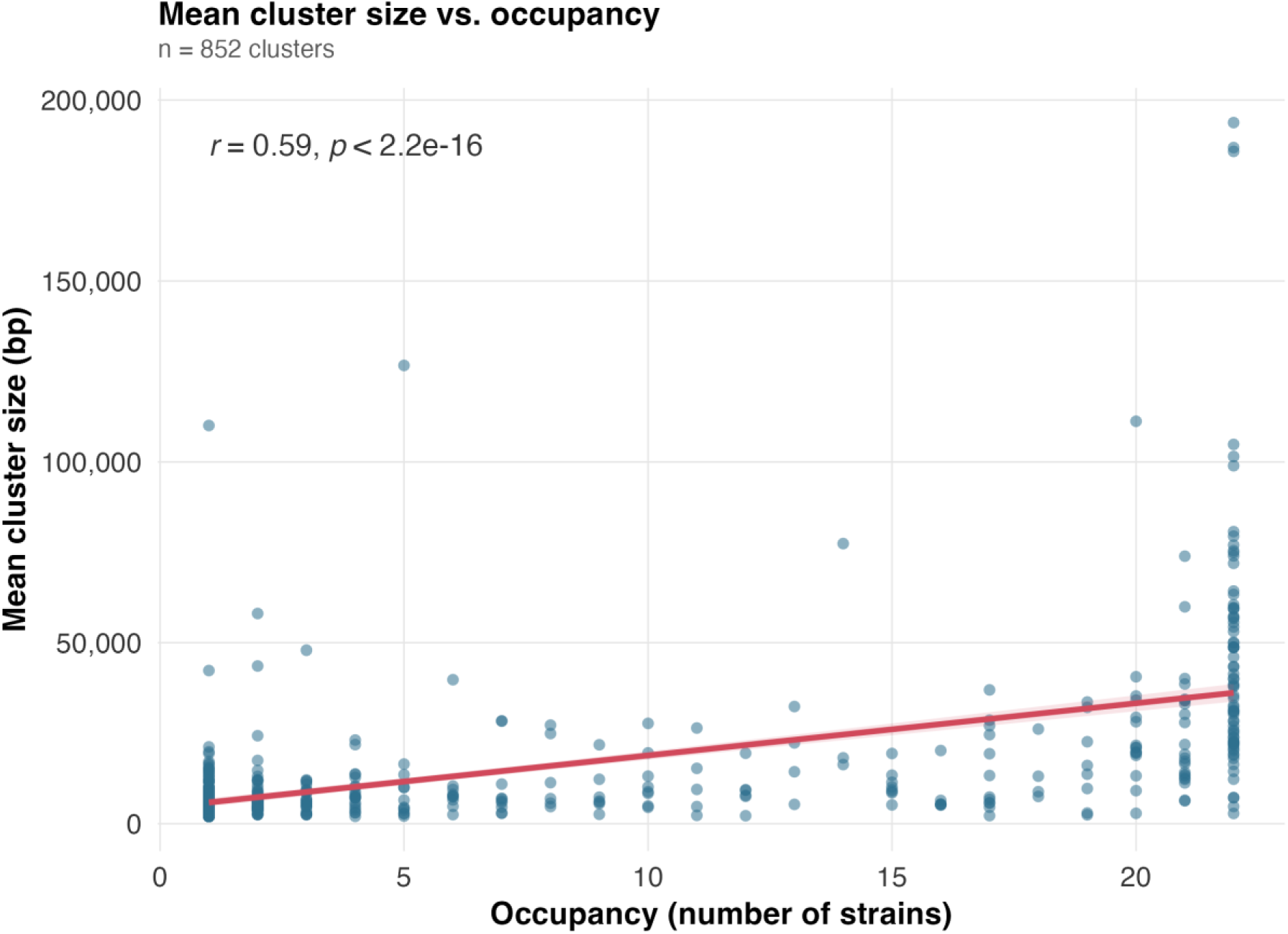
Relationship between piRNA cluster size and occupancy across strains. Each point represents one of the 852 non-redundant piRNA clusters identified across the 22 strains. Cluster size corresponds to the mean size of each cluster across strains in which it was detected. The red line shows the linear fit and the shaded area the 95% confidence interval. Cluster size was positively correlated with occupancy (Pearson’s r = 0.59, P < 2.2 × 10⁻¹⁶).

**Figure S10:**
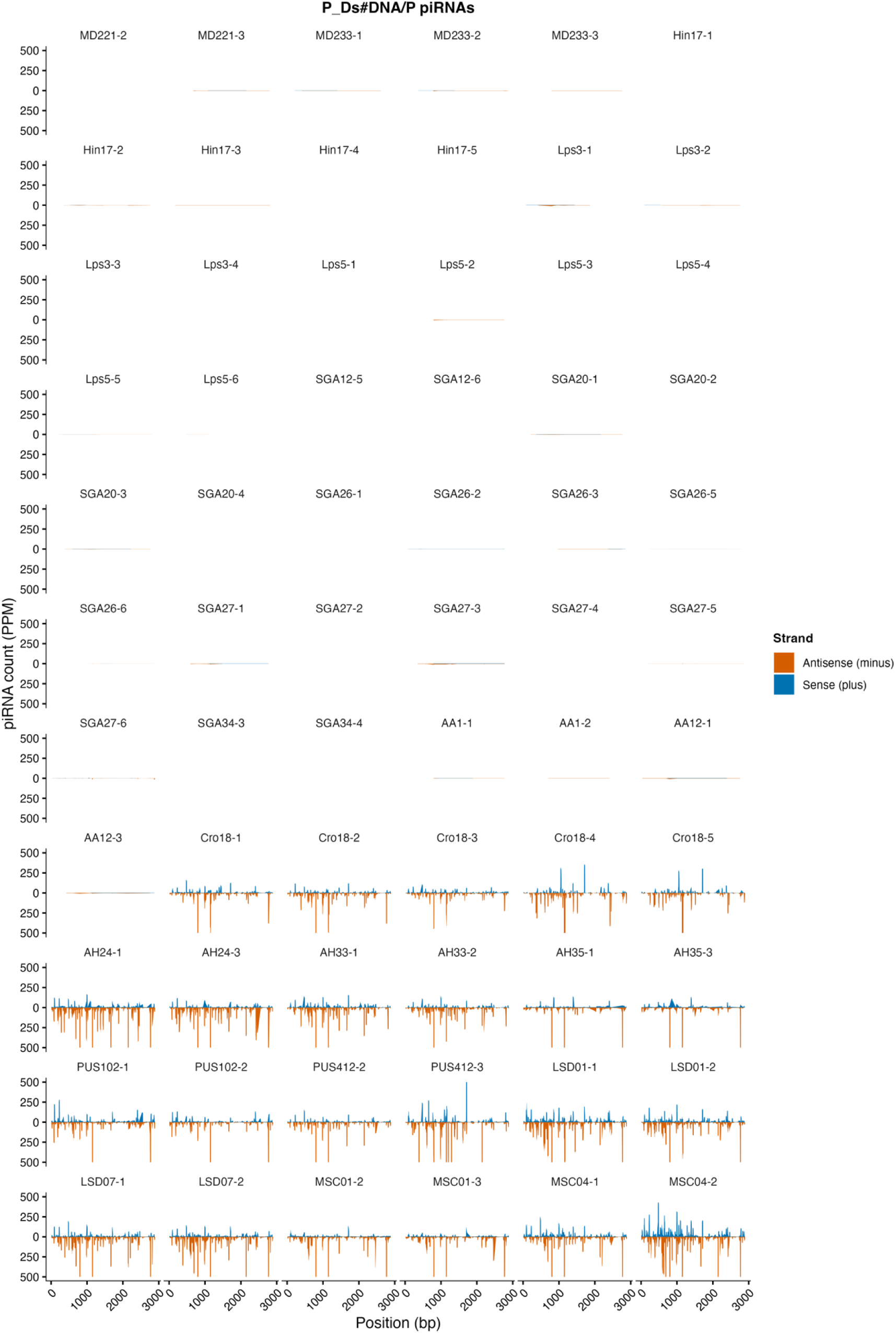
piRNA coverage profiles across the P-element consensus sequence for all replicates. Coverage is represented as PPM. To improve the visualisation, the y-axis is capped at 500 PPM.

**Figure S11:**
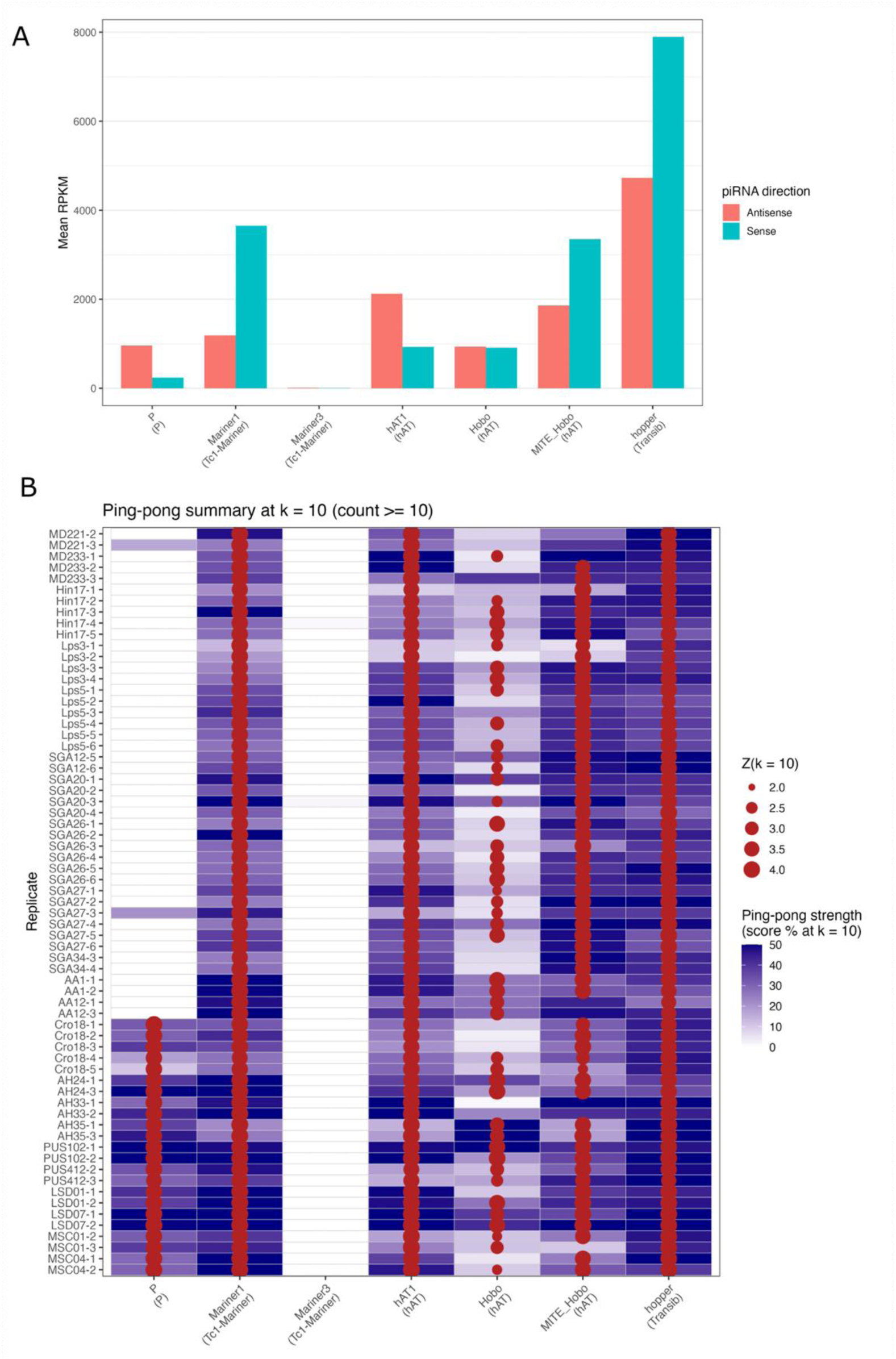
piRNA abundance and ping-pong signatures across selected DNA families in D. simulans. *(A) Mean sense and antisense piRNA abundance (RPKM - reads per kb per million mapped miRNA) for each TE family across all analysed replicates/strains. (B) Ping-pong signatures for each TE family across replicates/strains. Circle size represents the Z-score at the characteristic 10 nt overlap position (minimum Z = 2 and at least 10 overlapping read pairs), while the background colour indicates the relative ping-pong strength (% of overlaps at k = 10). TE superfamilies are indicated in brackets*.

**Figure S12:**
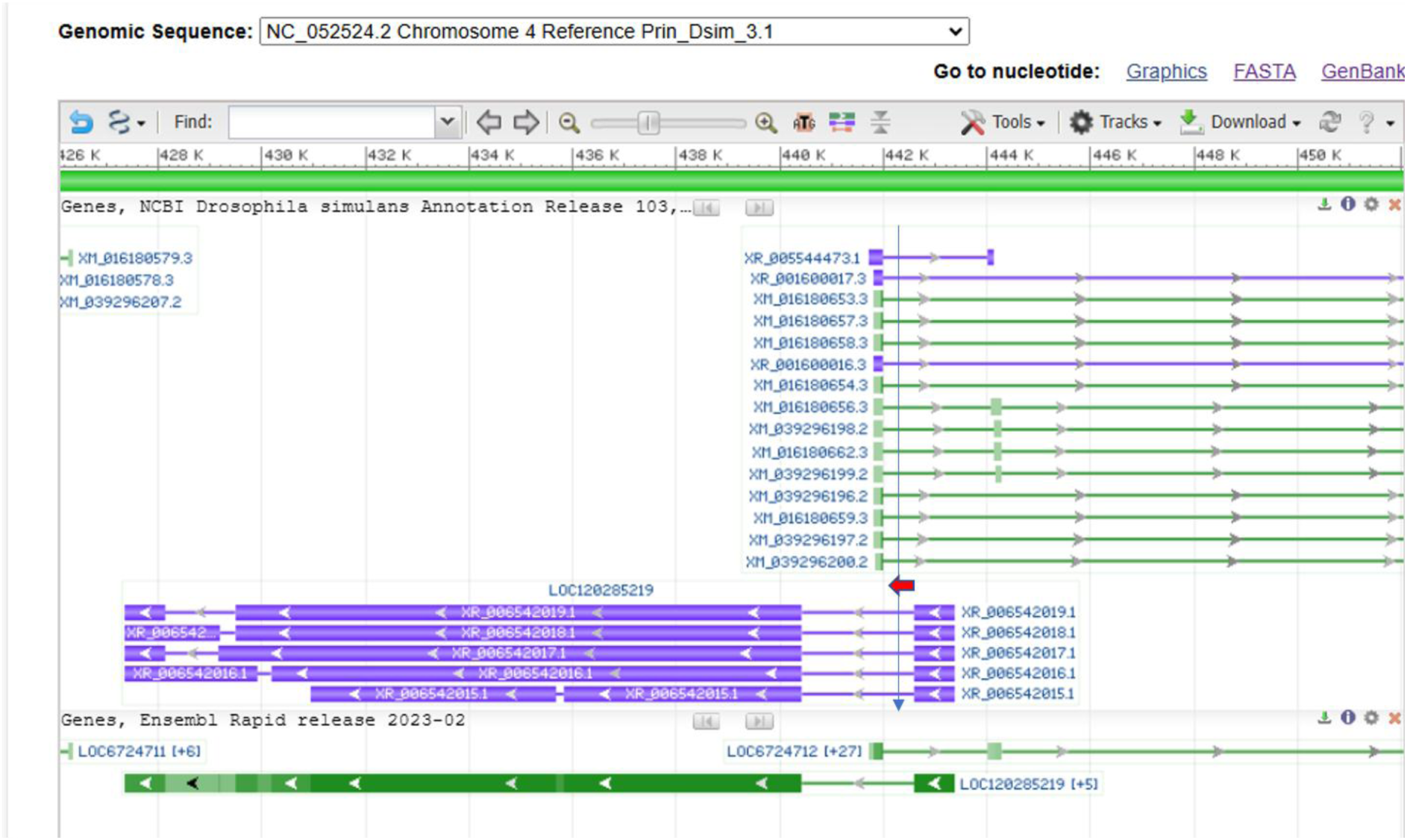
Screenshot from the NCBI Genome Data Viewer showing the reference genome annotation surrounding the genomic position of an exclusive P-element insertion identified in the Cro18 strain. The blue arrow marks the insertion site, and the red arrow indicates the direction of the P-insertion. The insertion is inserted in the antisense orientation relative to the protein-coding gene LOC6724712 and in the sense orientation relative to the overlapping uncharacterized lncRNA LOC120285219.

**Figure S13:**
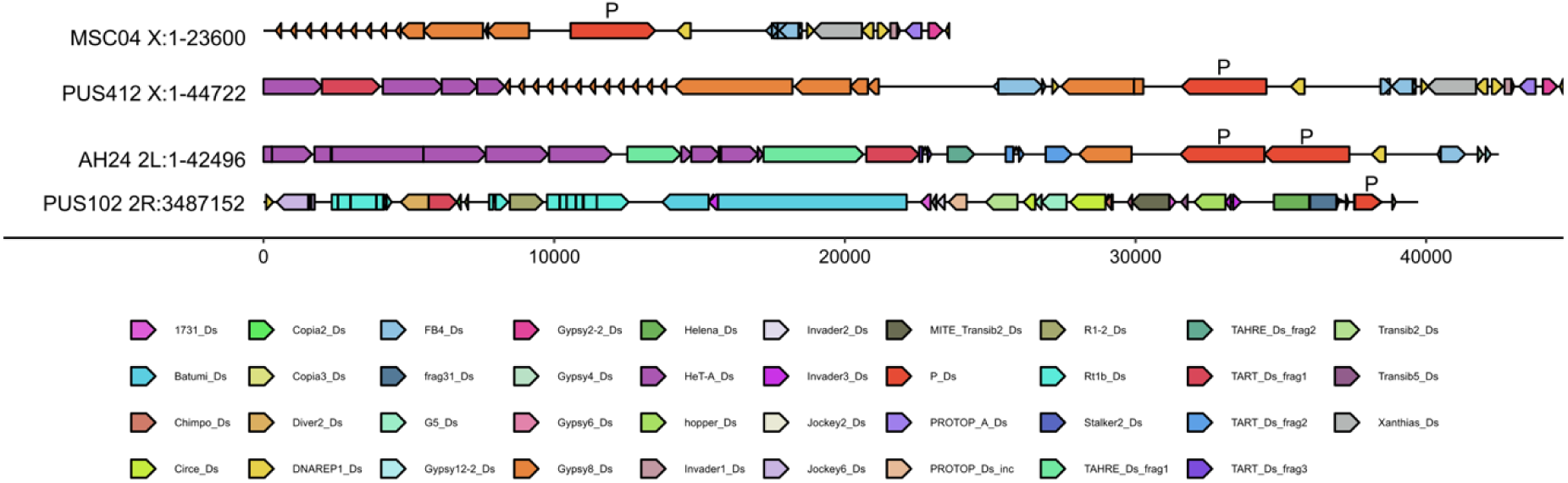
Illustration of piRNA clusters having the P-element, with colored segments indicating distinct TEs and orientation.

**Figure S14:**
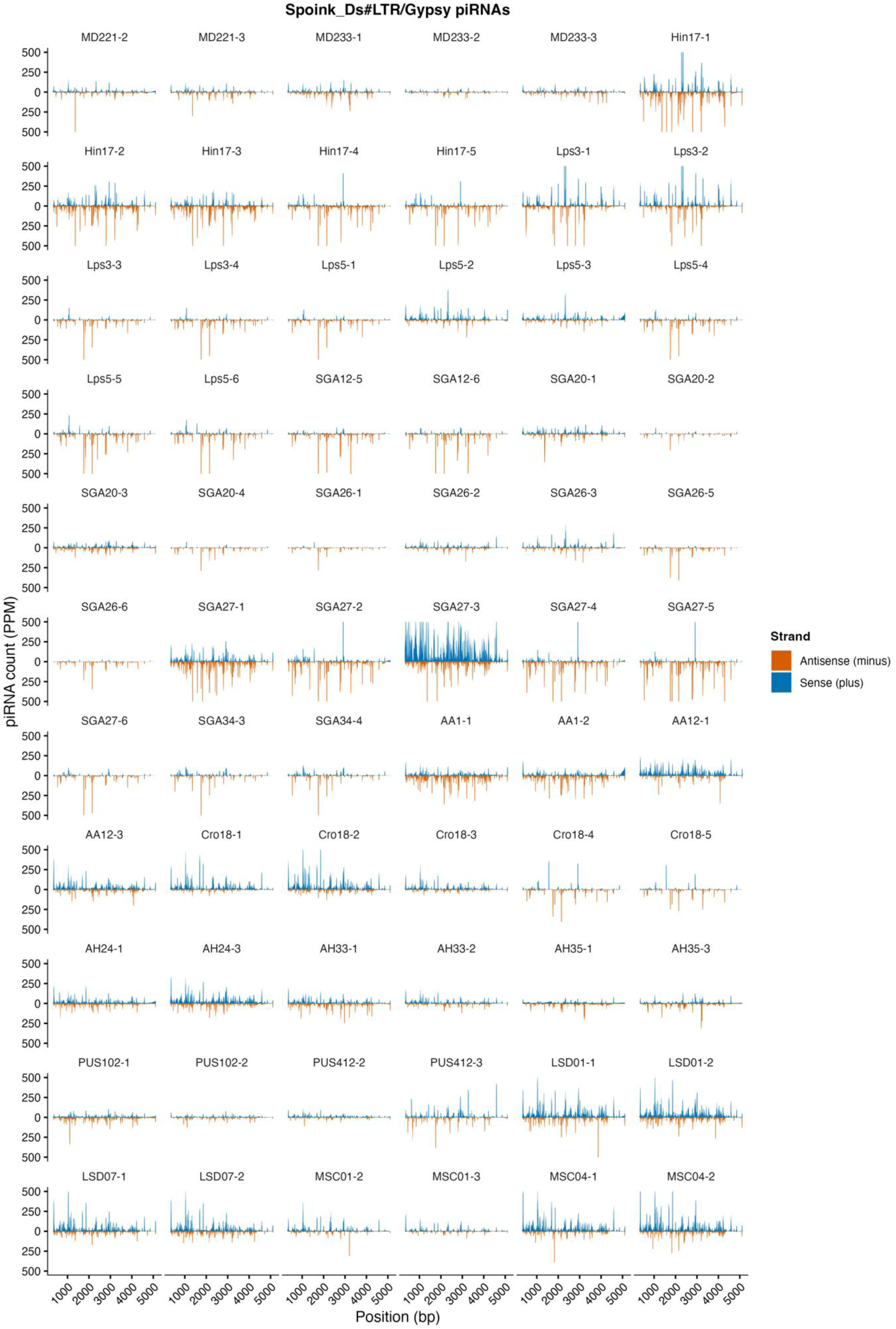
piRNA coverage profiles across the Spoink consensus sequence for all replicates. Coverage is represented as PPM. To improve the visualisation, the y-axis is capped at 500 PPM.

**Figure S15:**
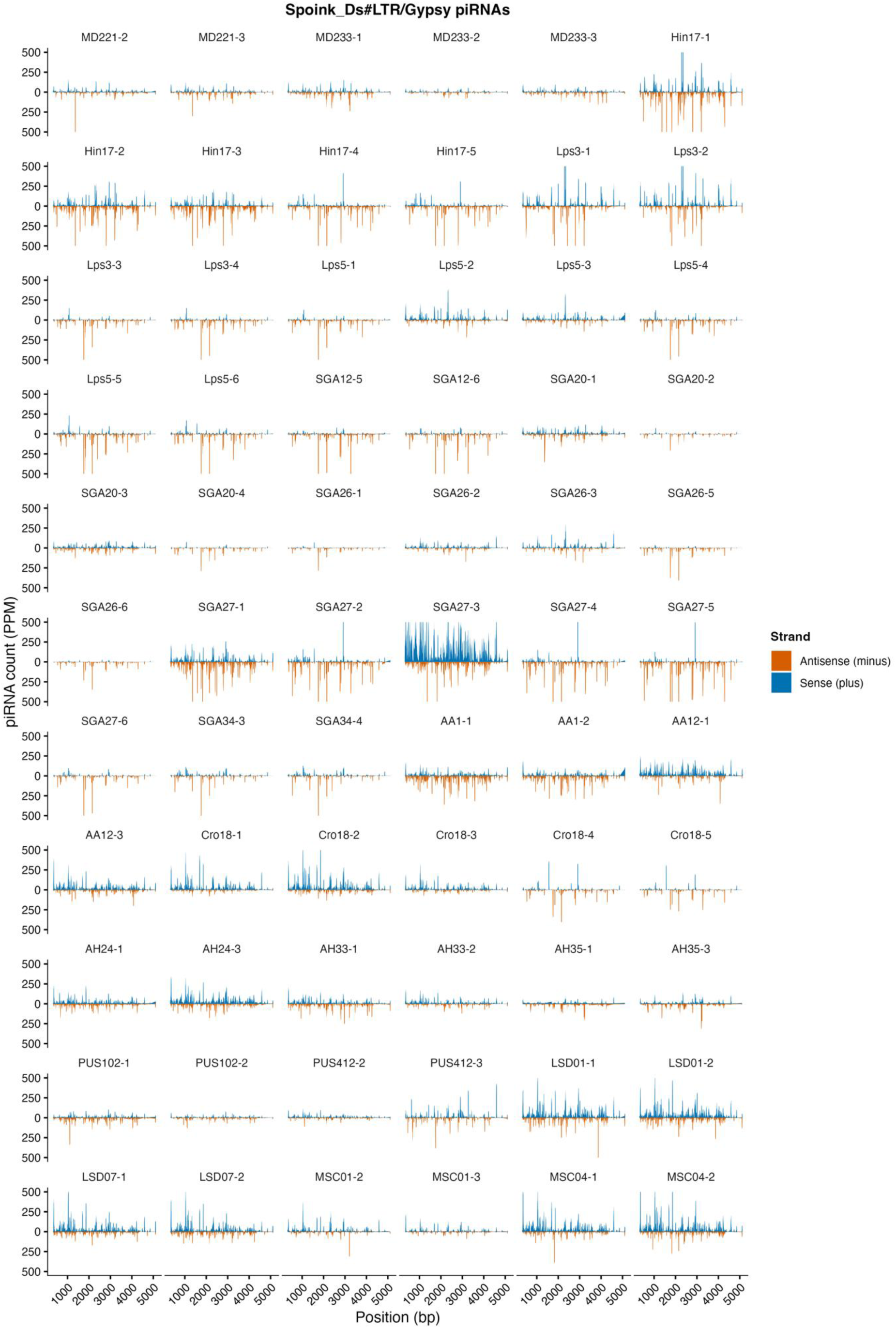
piRNA coverage profiles across the Shellder consensus sequence for all replicates. Coverage is represented as PPM. To improve the visualisation, the y-axis is capped at 500 PPM.

**Figure S16:**
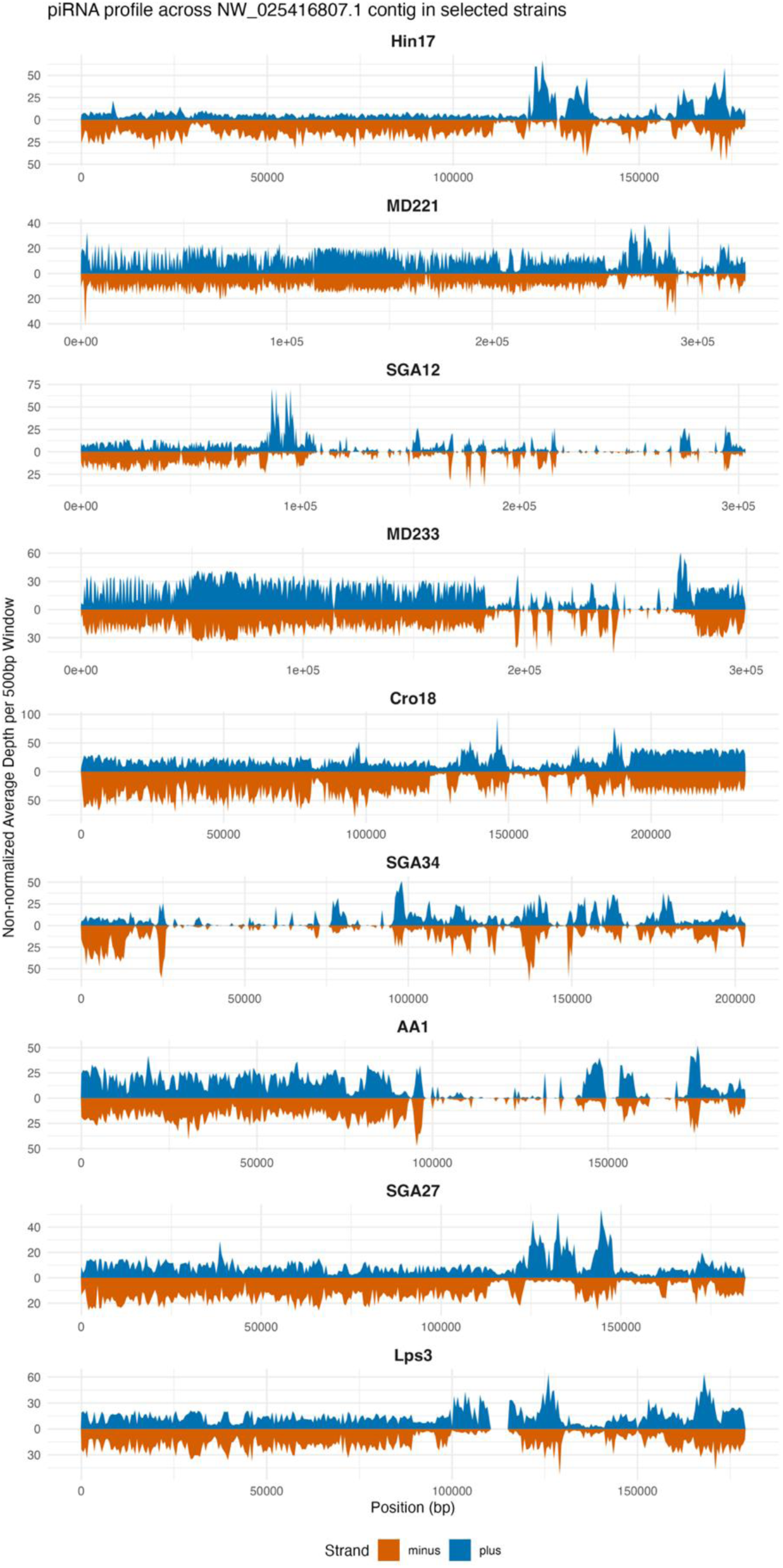
piRNA profiles across the centromeric satellite-containing contig (NW_025416807.1) in representative D. simulans strains. Blue and orange represent piRNAs mapping to the plus and minus strands, respectively.

**Figure S17:**
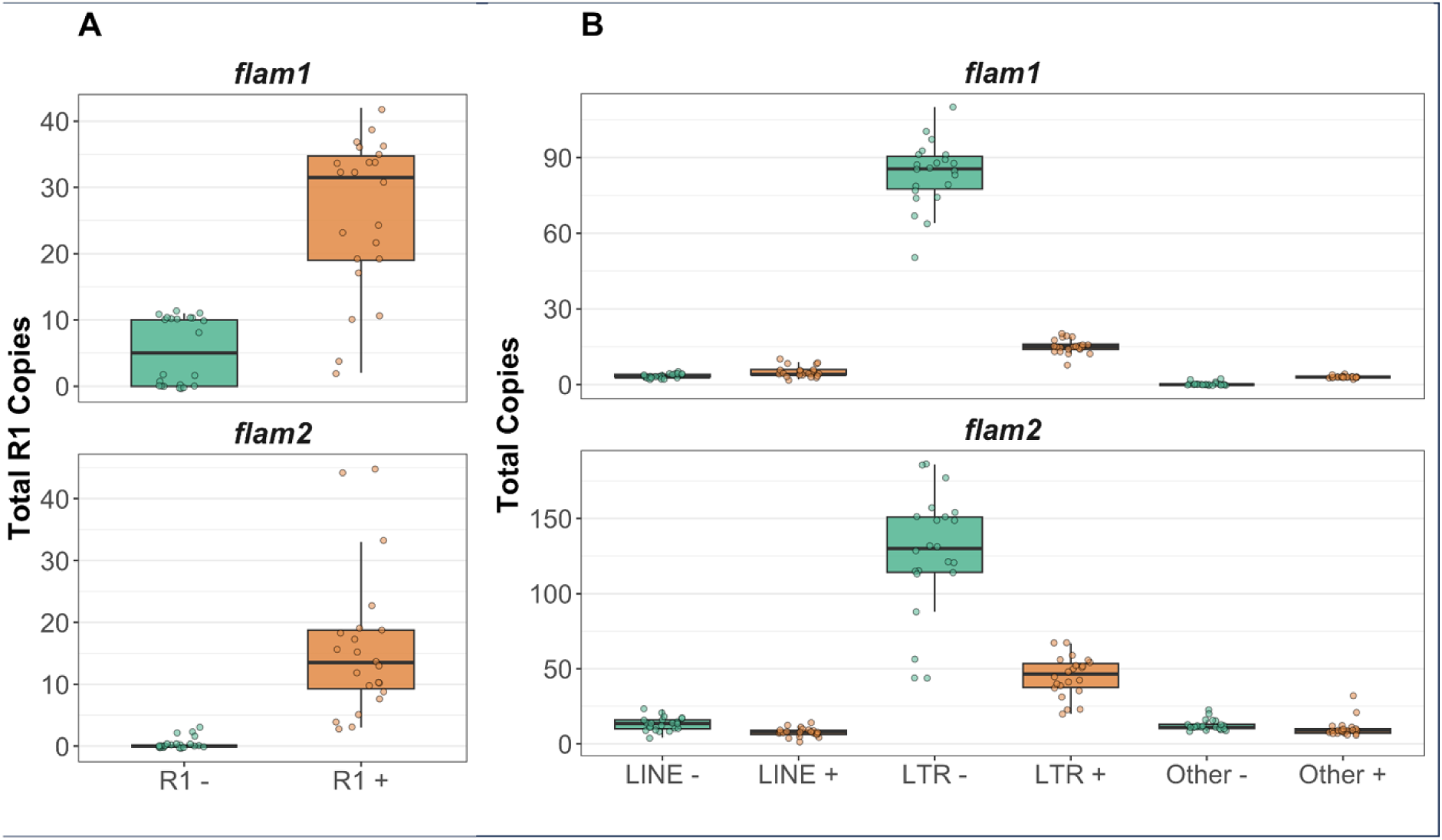
Composition of TEs in flam1 and flam2 across the 22 D. simulans genomes. (A) Total number of R1 copies; (B) Total copies of LINE, LTR and other TEs. (−, antisense; +, sense).

**Figure S18:**
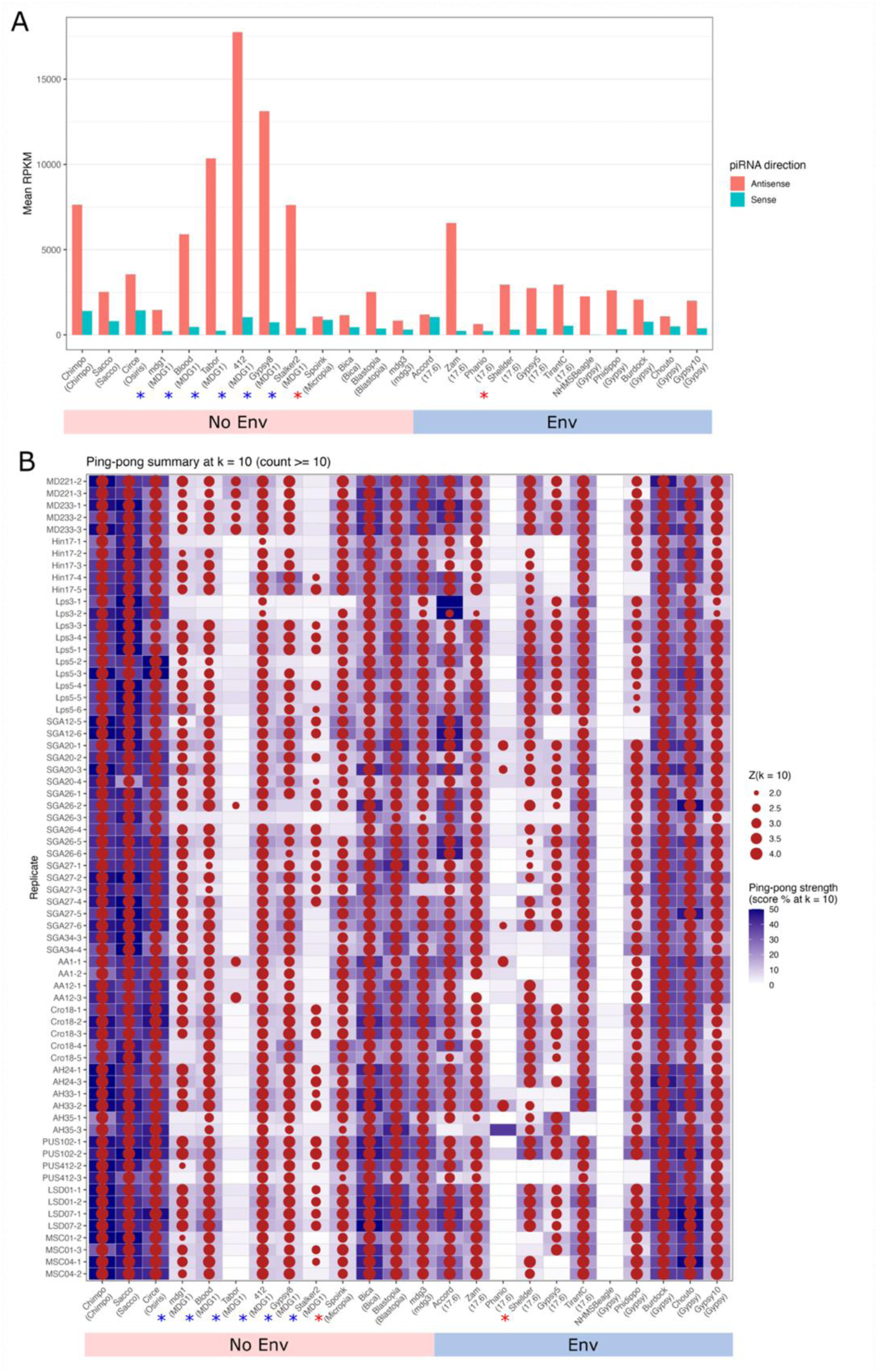
piRNA abundance and ping-pong signatures across selected LTR families in D. simulans. *(A) Mean sense and antisense piRNA abundance (RPKM - reads per kb per million mapped miRNA) for each TE family across all analysed replicates/strains. (B) Ping-pong signatures for each TE family across replicates/strains. Circle size represents the Z-score at the characteristic 10-nt overlap position (minimum Z = 2 and at least 10 overlapping read pairs), while the background colour indicates the relative ping-pong strength (% of overlaps at k = 10). LTR families are grouped by the presence or absence of env gene as identified during curation. LTR clades are indicated in brackets. Blue asterisks indicate LTR families belonging to the MDG1 clade, which lack an env gene but encode an accessory protein sORF2 that has been implicated in cell-to-cell transmission. Red asterisks highlight Spoink and Shellder*.

**Tables S1-S9** are available as Excel files at the following link.

https://zenodo.org/records/23987502?preview=1&token=eyJhbGciOiJIUzUxMiJ9.eyJpZCI6ImQyNTFiZmM5LTcwZGMtNGNmMS05YjZhLWFmM2Q5YzkyMjVmOCIsImRhdGEiOnt9LCJyYW5kb20iOiIyNWMyMjZhNzQxMjNkYWE5MDljOTM1YzcwMGIyYjY5ZiJ9.K4o9BVOJgrmnqP8Nw-usDr_Y_wU7ihzHwGXXsMT0QWwFTkk55db7WVr_3ga9IDFa8YUwYWSMj_nb7Z-DgdSErg

